# People with a tobacco use disorder misattribute non-drug cues as worse predictors of positive outcomes compared to drug cues

**DOI:** 10.1101/2023.03.27.534463

**Authors:** Shivam Kalhan, Philipp Schwartenbeck, Robert Hester, Marta I. Garrido

**Affiliations:** – The University of Melbourne, Melbourne School of Psychological Sciences, Melbourne, Victoria, Australia; – Graeme Clark Institute for Biomedical Engineering, Melbourne, Victoria, Australia; – University of Tübingen, Tübingen, Germany; – Max Planck Institute for Biological Cybernetics, Tübingen, Baden-Württemberg, Germany

## Abstract

Adaptive behaviours depend on dynamically updating internal representations of the world based on the ever-changing environmental contingencies. People with a substance use disorder (pSUD) show maladaptive behaviours with high persistence in drug-taking, despite severe negative consequences. We recently proposed a salience misattribution model for addiction (SMMA; Kalhan et al., (2021)), arguing that pSUD have aberrations in their updating processes where drug cues are misattributed as strong predictors of positive outcomes, but weaker predictors of negative outcomes. We also argue that conversely, non-drug cues are misattributed as weak predictors of positive outcomes, but stronger predictors of negative outcomes. However, these hypotheses need to be empirically tested. Here we used a multi-cue reversal learning task, with reversals in whether drug or non-drug cues are currently relevant in predicting the outcome (monetary win or loss). We show that compared to controls, people with a tobacco use disorder (pTUD), do form misaligned internal representations. We found that pTUD updated less towards learning the drug cue’s relevance in predicting a loss. Further, when neither drug nor non-drug cue predicted a win, pTUD updated more towards the drug cue being relevant predictors of that win. Our Bayesian belief updating model revealed that pTUD had a low estimated likelihood of non-drug cues being predictors of wins, compared to drug cues, which drove the misaligned updating. Overall, several hypotheses of the SMMA were supported, but not all. Our results implicate that strengthening the non-drug cue association with positive outcomes may help restore the misaligned internal representation in pTUD.

## 1. Introduction

In a non-random environment, the brain integrates past experience with current sensory input to make predictions about future environmental contingencies (Friston, 2010; Knill & Pouget, 2004; Körding & Wolpert, 2004; Tenenbaum et al., 2006). These predictions facilitate adaptive action selection. Therefore, one important aspect of brain functioning is to integrate past experience with the current sensory input such that the most accurate predictions are generated, which in turn leads to adaptive behaviors. *Internal representations* that best denote the environmental contingencies is key to making accurate predictions. However, to best represent the ever-changing environmental contingencies, these internal representations need to be dynamic, and constantly updated from new sensory information. Given that there is usually an abundance of sensory information, these internal representations also need to be selectively updated from information or cues that are most *relevant or salient* in predicting future outcomes. Internal representations are therefore most adaptive when they are selectively updated from cues with high predictive values (Mackintosh, 1975). And because these internal representations are used for action selection (Tolman, 1948), behaviors are only as adaptive as these internal representations.

Addiction is a complex condition often characterized by a high persistence in maladaptive behaviors, despite severe negative consequences (*Diagnostic and Statistical Manual of Mental Disorders (5th Ed.)*, 2013). We recently proposed a conceptual theory suggesting that aberrations in internal representation updating processes may be one contributor of some aspects of maladaptive addictive behaviors (Kalhan et al., 2021). Specifically, we proposed that drug predictive cues are misattributed as having a high salience in predicting positive outcomes, and less so in predicting negative outcomes. And the converse may also be true for non-drug predictive cues, with higher salience in predicting negative outcomes over positives. A consequence of this salience misattribution is that a misaligned internal representation is produced, where overweighted positive predictions/expectations in response to drug predictive cues are generated, thereby, increasing the chances of selecting drug-related actions. Further, the salience of any negative outcomes of drug actions may be downweighed, contributing to reduced updating of the internal representation from these drug-related negative outcomes. Therefore, a misaligned internal representation is formed where drug cues have an inaccurately higher predictive value for positive outcomes, and inaccurately lower for negative outcomes. We proposed that this misaligned internal representation may be one contributor in the high persistence of maladaptive drug-related behaviors, despite their negative consequences. We refer to our theory throughout the current paper as the salience misattribution model for addiction (SMMA theory).

The present experiment was designed to empirically test the theoretical concept of a salience misattribution effect producing a misaligned internal representation, between smoking-related (drug) and non-drug (neutral) cues in people with a tobacco use disorder (TUD) and non-smoking healthy controls. Here, we used a reversal learning paradigm, with reversals in the *relevance* of drug and neutral cues in predicting future outcomes, which could be a monetary win or loss. This was a non-instrumental task, where participants’ actions could not influence the outcome, and was adapted from Schwartenbeck et al., (2016) and Nour et al., (2018). Participants had to keep track of reversals but did not need to learn any new cue-outcome relationships during the experiment.

We used a simple trial-by-trial Bayesian belief updating model (see *Methods, computational modelling*) to estimate how much predictive value a participant placed on drug and neutral cues (all of which had objectively equal predictive values in the task). An internal representation in this task is specifically defined as a representation of whether the neutral or drug cue type is currently relevant in predicting the outcomes. The Bayesian model worked by integrating past experience (*prior belief* of which cue type is currently relevant), with the incoming sensory information (*likelihood* of the current outcome, given the cues presented), to produce an updated belief of which cue type is currently relevant (*posterior belief*). Using this Bayesian model, we estimated a trial-by-trial belief estimate of whether the drug or neutral cues are currently relevant in predicting the outcome. We also calculated the Kullback-Leibler divergence (KLD) on a trial-by-trial basis, which is a difference between the prior and posterior belief distributions and quantifies the magnitude of internal representation updates (information gain). Lastly, we used eye-tracking (gaze proportions and pupil size) to test whether eye-behavior correlated with internal representation updating.

Overall, we hypothesized that for people with a TUD, more so than in controls: 1) drug cues, compared to neutral cues, would have a higher estimated predictive value for positive outcomes, and 2) neutral cues, compared to drug cues would have a higher estimated predictive value for negative outcomes. The differences in predictive values may therefore capture a salience misattribution effect, that produces misaligned internal representations of cue-outcome predictions in people with a TUD, due to aberrant updating.

## 2. METHODS

### 2.1. Participants

Participants were recruited through advertisements at the University of Melbourne, paid Facebook advertisements, and a community website (gumtree.com.au). All participants provided written and verbal informed consent that was approved by the Human Ethics Committee of the University of Melbourne. The TUD group consisted of 24 individuals (12 males, 12 females: mean age 34.54, standard deviation +/- 9.96, and range 19-53). People in the TUD group smoked at least 10 cigarettes daily, for at least the last 2 years. The average Fagerström Test for Nicotine Dependence (FTND) (Heatherton et al., 1991) score was 4.92, indicating moderate dependence. The control group consisted of 24 individuals (12 males, 12 females; mean age 34.33, standard deviation 9.76, and range 18-55). Participants in the control group reported smoking fewer than 5 cigarettes in their entire lifetime and having no history of vaping or neurological/psychiatric disorders. The mean Alcohol Use Disorders Identification Test (AUDIT) (Saunders et al., 1993) score for the control group was 2.96 with standard deviation +/- 3.69, and 10.38 with standard deviation +/- 9.04 for the TUD group. All participants were paid $20 per hour for their time during the experiment, and the TUD group participants were paid an additional $30 for abstaining 3-hours before the study.

### 2.2. Procedure

People in the TUD group were asked to abstain from smoking at least 3-hours prior to participating in the experiment, which was confirmed by self-report and further supported by the mean Shiffman–Jarvik Withdrawal Scale (SJWS) (15 items) (Shiffman & Jarvik, 1976) score of 4.13 +/ 0.65, indicating moderate craving/withdrawal symptoms. The 3-hours abstinence window was chosen due to nicotine’s half-life of approx. 2-hours (Benowitz et al., 1982), and data suggesting that the 3-hour abstinence did not produce withdrawal effects on cognition (Charles-Walsh et al., 2014). At the start of the experimental session, participants were asked to fill out the relevant questionnaires. For the control group, this was the AUDIT questionnaire and for the TUD group this was the FTND and SJWS, in addition to the AUDIT. Next, participants from both groups were asked to rate all four visual cues used in the task for valence, arousal, and the urge to smoke. These cues were rated on a computerized slidable scale between 0 and 100, which was similar to how it was used by Manoliu et al., (2021) to assess cue ratings in people with a TUD. Subsequently, participants were given written and verbal instructions on the task, which took approx. 10 minutes. Next, participants did the training task (approx. 10 minutes – see *Training session* below). After this training task, we performed the eye calibration procedure (usually took less than 2 minutes). And finally, the participant did the main experiment (approx. 45 minutes), with short breaks every ∼12 minutes.

### 2.3. Task contingencies and trial types

The current experiment used a reversal learning task with reversals in the *relevance* of the cue type (drug or neutral) for predicting future monetary outcomes (wins or losses). Sometimes drug cues were relevant in predicting the future outcomes, and this would then reverse to neutral cues being relevant in predicting the outcome. At the beginning of every trial, one drug and one neutral cue were visually presented side-by-side. The participant’s primary aim was to keep track of these reversals and indicate, on a trial-by-trial basis, *which cue type they thought was currently relevant in predicting the outcomes*. They also indicated, on a trial-by-trial basis, how confident they were in their decision. Importantly, to minimize any explicit bias, we did not directly ask participants if they thought the “drug” or “neutral” cue predicted the outcome. Instead, participants were therefore asked whether they thought the “circle” or the “square” predicted the outcome, as both (drug and neutral) cues were in either a circle or a square outline. In other words, if drug cues were consistently presented in a circle, neutral cues would be consistently presented in a square throughout the entire task for that participant. The outline for the drug and neutral cues was either a circle or a square and was counterbalanced across participants.

There were four visual cues used in total - two drug cues and two neutral cues. One drug and one neutral cue predicted a win 80% of the time, and a loss the other 20% of the time. The other drug and neutral cue predicted a loss 80% of the time and a win the other 20% of the time (see Figure 1a for an example). The specific drug and neutral cues that predominately predicted wins/losses were counterbalanced across participants. Participants could infer which cue type was currently relevant in predicting the outcome when the two cues that were presented side-by-side, predicted opposite outcomes (*informative trials*). For example, if the predominately win predictive drug cue and the predominately loss predictive neutral cue were presented side-by-side, and the outcome was a win – it was likely that the drug cue predicted this, and the participants could therefore infer that they are likely in the drug relevant state (Figure 1b). These trials, where the two cues predicted opposite outcomes (incongruent trials; i.e., presenting one win predictive cue and one loss predictive cue) were informative trials as these could help the participant infer which cue predicted the outcome, indicating whether they were in a drug or neutral cue relevant state. These informative trials made up 40% of the total trials. Another 40% of the trials were *uninformative*, where both cues predicted the same outcome (congruent trials; i.e., both cues were win or loss predictive). These trials were not informative because they did not allow the participant to infer which cue predicted this outcome, and therefore updating/changing their estimate of which cue type was relevant would mean that they were using unreliable information to guide their inferences.

**Figure 1.**
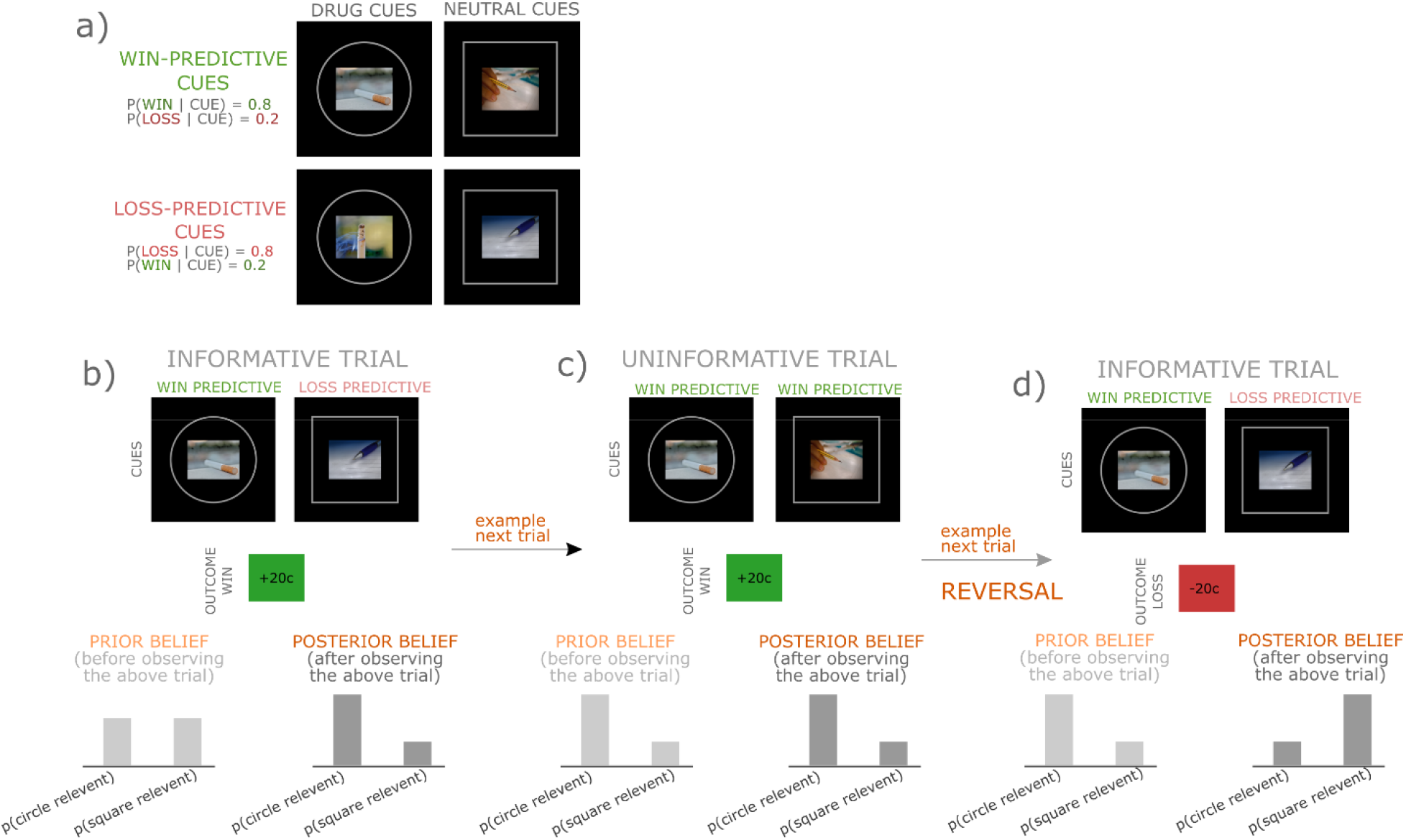
Experimental contingencies and trial types. a) An example of cue contingencies for a given participant. There is always one drug and one neutral cue predominately predictive of a win outcome (p(win | win cue) = 0.8, p(loss | win cue) = 0.2), and another drug and neutral cue predominately predictive of a loss outcome (p(loss | loss cue) = 0.8, p(win | loss cue) = 0.2). Drug cues are inside circles and neutral cues inside squares in this example. b) Example of an informative trial. The drug cue (circle) predicts a win, and the neutral cue (square) predicts a loss. As the outcome is a win, the participant can infer that they are likely in the drug relevant state (circles). Bottom panel is a situation where the participant had the prior belief (belief before the trial) that both drug and neutral cue were equally relevant in predicting the outcome. Then the posterior belief (belief after observing that trial) may be updated, to being more likely that the participant is in a drug relevant state, and less likely in the neutral state. C) Example of an uninformative trial. Both the drug and the neutral cues are predictive of wins, and the outcome is a win. Therefore, this is not informative about which cue type is currently relevant. The bottom panel shows no change in the posterior belief after observing an uninformative trial. A change in the posterior belief would indicate that the participant is using irrelevant information to make their inferences. D) Another informative trial, but indicative of a reversal. Previously, the drug cues were predictive of the outcome. In this case the drug cue predicts a win outcome, and neutral cue predicts a loss outcome. Because the outcome is a loss, the participant could now infer that the neutral cue (square) is likely now the relevant cue type. Bottom panel shows the prior belief of the drug cue (circle) being relevant, but after observing the trial, which indicates a reversal has occurred, the updated posterior belief is now indicative of the neutral cue (squares) being the relevant cue type.

The last 20% were *surprise trials,* 10% of these were incongruent surprises, when participants were in a drug or neutral relevant state and the outcome was opposite to what the drug or the neutral cue would have predicted (e.g., if the participant was in a drug relevant state and the drug cue was predictive of a win, but the outcome was a loss) and the other 10% were congruent surprises where the outcome would be the opposite to what both cues predicted (e.g., the drug and neutral cues predicted a win, but the outcome is a loss). The distribution of these surprise trials were constrained such that there were no repeats, and the very first trial could not be a surprise trial. There were 280 trials in total, 112 were informative (40%), 112 were uninformative (40%), 28 were incongruent surprises (10%) and 28 were congruent surprises (10%). See Table 1 for a summary of the trial types and the relative proportions.

The relevance of the cue type in predicting the outcomes reversed 24 times in total, with a reversal every 8, 10, or 12 trials. This number of trials before a reversal occurred was an even number so that there was an equal number of informative and uninformative trials within a relevance block. Whether the first relevant cue state was drug or neutral was counterbalanced between participants. Lastly, there was at least one informative trial within the first three trials after the reversal. This allowed the participant to infer that a reversal had occurred.

Participants were instructed that each cue had 80% predictive values, and that 20% of the time they would predict the opposite outcome. They were also instructed that there would be a reversal every 8-12 trials. They were not informed on the proportion of informative vs uninformative trials. Participants were also trained on which drug and neutral cue was predominately win/loss predictive before starting the task (see the *Training Session* section below). Importantly, while the money won or lost was real money, their responses did not influence these win/loss outcomes. These outcomes were predetermined.

### 2.4. Experimental Paradigm

The task started with a fixation screen randomized to be between 0.5-0.75 s long, following which the two cues were presented side-by-side for 2 s (see Figure 2; cue epoch). One was a drug cue, and the other was a neutral cue. The side (left or right) of cue presentation (neutral or drug) was randomized between trials. Following this 2 s cue epoch there is a blank ‘anticipation’ screen presented for 1.5 s. After this, the participant is presented with the outcome for 2 s. If it’s a loss, the square is red, and the sign is negative (“-”) with the amount lost (randomized between 15 and 20 cents) indicated in cents with the “c” symbol. In the case of a win, the square is a green and the sign is positive (“+”) with the amount won (randomized between 20 and 25 cents) indicated in cents. While there were an equal number of trials with a win and loss, wins had a larger amount than losses so that participants could gain money on average at the end of the task. All participants went through the same win-loss contingencies – with $4.89 received from the task itself, by all participants (this, in addition to the money received for their time – see *Participants* section above).

**Figure 2.**
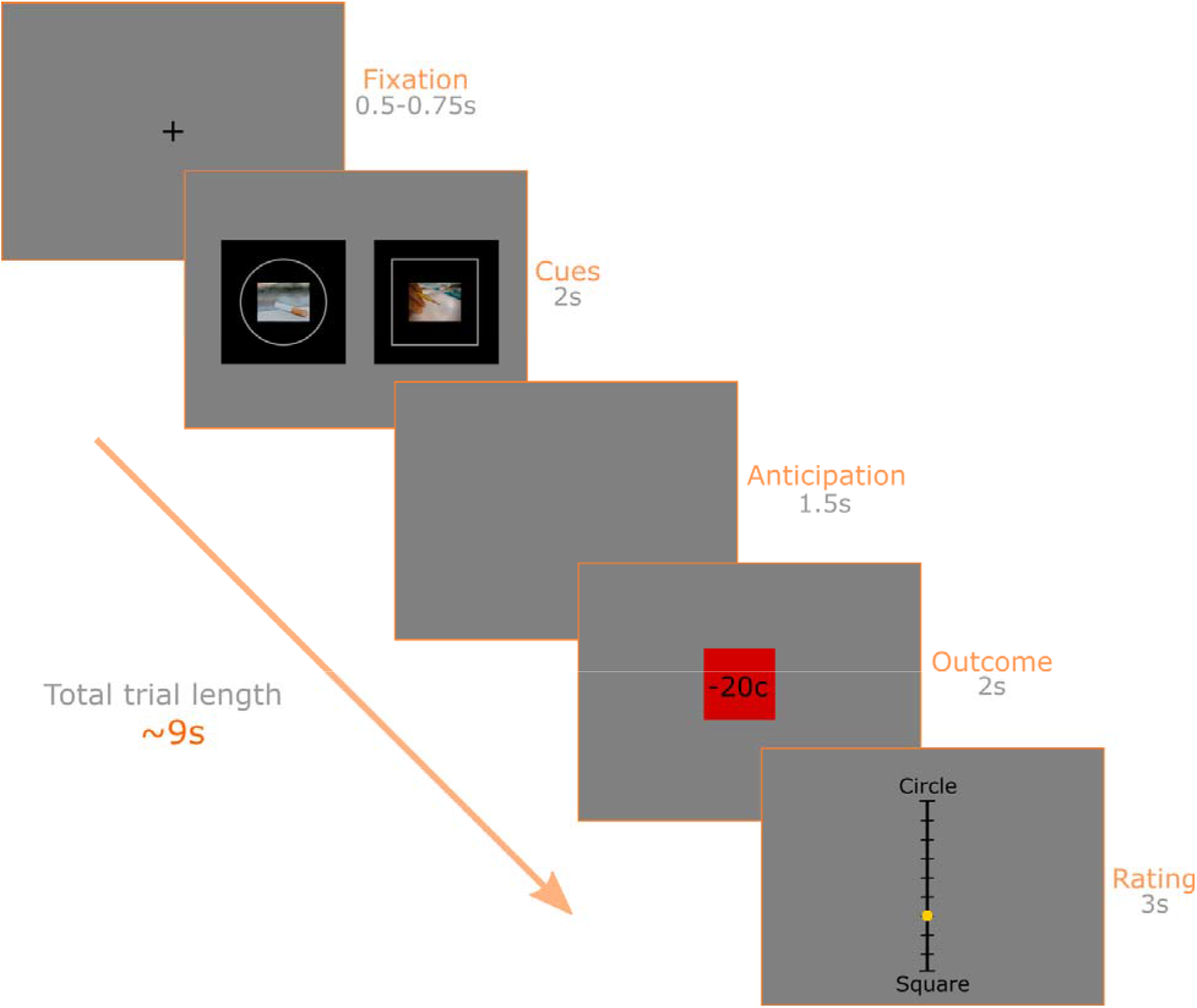
Example of an experimental trial. The trial starts with a fixation screen which is randomised to be between 0.5-0.75 seconds (s) long. Following this, the drug and the neutral cues are presented side-by-side for 2 s. Then a blank ‘anticipation’ screen appears for 1.5 s. After this blank screen, the participant gets the outcome for 2 s which, in this case, is a loss. The amount lost is indicated in cents. Lastly, a rating scale is presented. The participant has 3 s to rate how confident they are about whether they are in the circle (i.e., drug) or the square (i.e., neutral) relevant state, by moving the yellow dot up or down. The position of the yellow dot is recorded as their response at the end of the 3 s.

Finally, participants were presented with a vertical rating scale (11 points; numbers not indicated in the scale), where they could select whether they thought that the circle or the square predicted the outcome. We used a vertical scale, instead of horizontal to avoid a left-right bias based on the way the cues were presented. Participants were instructed that this was a confidence scale, and to utilize the full scale (i.e., go all the way towards the “circle” end if they are very confident that the circle predicted the outcome, and vice versa towards the “square” end, or the middle if they’re completely unsure). The yellow dot indicated what the choice was on the scale, and this dot was randomly placed at the extreme end of either the circle or the square at the start of each trial. Participants therefore had to move the dot, using the “up” or “down” arrow on a keyboard, to indicate their choice. Participants had 3 s to make their rating choice, and after the 3 s, their choice was recorded, and the trial ended. Whether the “circle” text position or the “square” test position was at the top or bottom of the scale was counterbalanced between participants. We used COGENT on MATLAB 2017b to run this experiment, on a 1920 × 1080p monitor, with a 120Hz refresh rate.

### 2.5. Training session

The primary aim of the training session was to ensure that participants learnt which drug and neutral cues were predictive of a win or a loss. Unlike in the main task, all trials were informative (with 100% predictive values for all cues). There were no uninformative and no surprise trials. In the first trial, the participants were told which shape was relevant so that they could get started on the task. The timing of a trial and all the events within the trial were identical to the main task, with the participants having to rate the shape that they thought predicted the outcome. All 4 cues had 30 trials each of predicting the outcome, allowing for an equal opportunity to learn the contingencies across the 120 trials. All participants had learnt what outcome each of the four cues were predictive of prior to starting the main task (based on a positive correlation between their ratings and true task contingencies).

### 2.6. Stimuli

The two drug cues were chosen from the “SmoCuDa” database, which is an open access database of visual smoking cues validated to induce craving in people with a tobacco use disorder (Manoliu et al., 2021). They were chosen based on having similar valence (51 +/- 15. and 55 +/- 21), arousal (56 +/- 21 and 66 +/- 20) and urge to smoke (54 +/- 23 and 57 +/- 26) ratings. The neutral images were of a pen and pencil, which are commonly used as control images when comparing with smoking images (Kang et al., 2012; Valsecchi & Codispoti, 2022). These images were downloaded from pixabay.com, a royalty free database of images. To make the images suitable for eye-tracking, all images were matched for luminance, spatial frequency, and the pixel histograms (contrast) using the “shine_color” toolbox (github.com/roddalben/shine_color) in MATLAB 2017b. These were all done using the “lumMatch”, “sfMatch” and “histMatch” functions within the toolbox. All four images were 209.55mm (width) × 157.16mm (height). We put a 500 × 500mm black square behind all these images. The grey (RGB: 153 153 153) outline was used to indicate the shape, and this was 350mm × 350mm. All images were in .png format, and were then converted to .bmp format, using the “imwrite” function in MATLAB 2017b in order to be used within COGENT. Overall, visual salience was very closely matched across all cues, which is considered best practice for eye tracking studies (Carter & Luke, 2020).

Lastly, the color green is usually responsible for a higher luminance and can have a greater perceived brightness compared to the color red (Cohen et al., 1968; Mullen, 1985). Our outcome squares (even though they were very small relative to the screen) were red or green for losses and wins, respectively. Therefore, we used the luminous efficiency function (Cohen et al., 1968) (Y = 0.21*Red + 0.72*Green + 0.07*Blue) to weight the red and green colors such that they had equal perceived brightness and minimized any confound on our pupillometry results due to the difference in colors.

### 2.7. Eye-tracking and calibration

The calibration was done prior to starting the main experiment and was a 10-point setting using the Eyelink toolbox for MATLAB (Cornelissen et al., 2002), which also used Psychtoolbox. Only the right eye was recorded. One eye is often recorded and has close accuracy and systemic error (sometimes lower) compared to two eyes which are later averaged (Carter & Luke, 2020; Hooge et al., 2019). We used the Eyelink® 1000 Plus eye tracker (https://www.sr-research.com/wp-content/uploads/2018/01/EyeLink-1000-Plus-Brochure.pdf) with 1000 Hz sampling frequency. There were 3 participants in the TUD group, and 4 participants in the control group excluded from eye-tracking analysis (but were still included in modelling and behavioral analysis), due to difficulties in tracking their eyes. Therefore, the eye-tracking data is based on n = 21 for the TUD group and n = 20 for the control group.

### 2.8. Computational Modelling

The current model was built within the Bayesian framework, using a Hidden Markov Model. The model was used to infer the current hidden relevant state (X) in the task, which could either be a drug or a neutral relevant state (*X* ∈ {1,2}, 1 = *drug relevent*, 2 = *neutral relevent*). The hidden relevant state is inferred based on three observations in a given trial: 1) the drug cue (D) presented (whether it was predominately predictive of a win or a loss; *D* ∈ {1,2}, 1 = *win predictive*, 2 = *loss predictive*), 2) the neutral cue (N) presented (whether it was predominately win or loss predictive; *N* ∈ {1,2}, 1 = *win predictive*, 2 = *loss predictive*)), and 3) the outcome (Y) (whether it was a win or a loss; *Y* ∈ {1,2}, 1 = *win*, 2 = *loss*)). These collectively formed the observation matrix (O), per every trial (t):*O_t_* = [*D_t_, N_t_, Y_t_*]. Given this observation matrix, the Bayesian model inferred the hidden relevant state on a trial-by-trial basis, where t denotes the current trial (*X_t_* ∈ {1,2}).

The model started with a uniform *prior belief,* which is a probability estimate of which hidden state is currently relevant under each of the two hypotheses (i.e., hypothesis one (H1) - the probability that the drug cues are currently relevant (*X* = 1) and, hypothesis two (H2) - the probability that neutral cues are currently relevant (*X* = 2)). These prior beliefs are weighted by the *likelihood* of observing the outcome given the cues that were observed during the trial, under each of the two hypotheses (see Equations 2 and 3 below). For example, if a win predictive drug cue and a loss predictive neutral cue was presented, and the outcome was a win, there is a higher likelihood that the drug cue predicted this win outcome. Therefore, the prior belief would be updated towards the drug relevant state (H1; X = 1). These prior beliefs and the likelihoods are used to calculate the *posterior belief*. This posterior belief is the updated probability estimate of the current hidden relevant state under both hypotheses, after observing the current trial. The posterior belief can be calculated on a trial-by-trial basis using Bayes’ rule below:

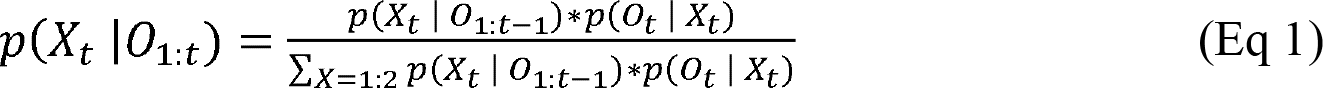

Where, *p*(*X_t_* | *O*_1:*t*_) is the posterior belief after the observations of the current trial (t), the *p*(*X_t_* | *O*_1:*t*-1_) is the prior belief, and *p*(*O_t_* | *X_t_*) is the likelihood of the observations under both hypotheses. Importantly, the prior belief of the next trial (t+1) is the posterior belief of the current trial (t) (see bottom panel of Figure 1b-d).

For every trial, the likelihood is based on the cues observed, for both hypotheses, based on the matrices see below:

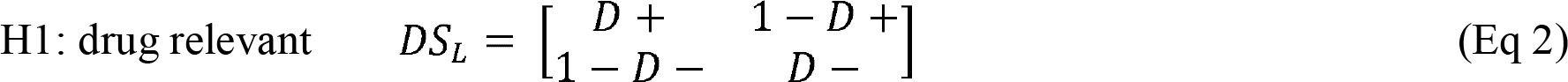

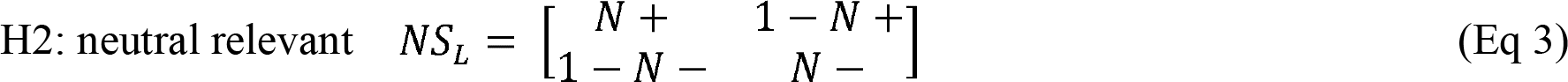

Here, *DS_L_* is the likelihood matrix under the hypothesis that the current hidden state is drug relevant (H1; X = 1), given the observations. The *NS_L_* is the likelihood matrix under the hypothesis that the current hidden state is neutral relevant (H2; X = 2), given the observations. Here, a win predictive drug and neutral cue predict the outcome win with likelihood D+ and N+, respectively. And a loss with likelihood 1-D+ and 1-N+, respectively. Conversely, a loss predictive drug and neutral cue predict a loss with likelihood D- and N-, respectively. And a win with likelihood 1-D- and 1-N-, respectively. Therefore, in both these matrices, row one represents a win outcome, and row two a loss outcome (*Y* ∈ {1,2}; 1 = *win*, 2 = *loss*). And in both these matrices, column one represents the win predictive cue (D+ and N+) and column two is the loss predictive cue (D- and N-) (*D and N* ∈ {1,2}; 1 = *win predictive*, 2 = *loss predictive*). Under this layout, the likelihoods can be chosen for the two hypotheses as below:

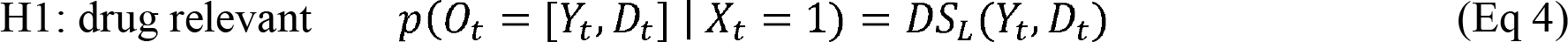

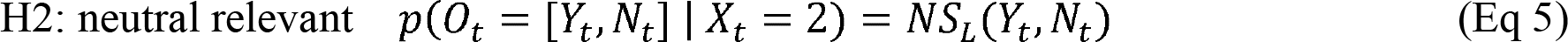

These four likelihood values (one for each cue) were free parameters and was estimated for each participant. The greater the likelihood estimation for a given cue, the more the estimated predictive value (likelihood) for that cue, and the more the belief updates based on observing this cue. The likelihood estimate could be between 0.5 and 1. A likelihood value of 0.5 means no predictive value (no updates based on this) and 1 means perfect predictive value (maximal belief updating). Importantly, the overall internal representation update depends on the likelihood of both hypotheses, where if the likelihood of H1 is much greater than the likelihood for H2, the update will be strongly in favour of H1. But if the likelihoods for the two hypotheses are similar, there will be a weaker update, in favour of the hypothesis with a higher likelihood. For example, if a drug positive cue was presented, and a neutral negative cue was presented, and the outcome was a win, the internal representation update is dependent on 1) the likelihood of the drug positive cue (D+) predicting this win outcome, and 2) the likelihood of the neutral negative cue (1-N-) predicting this win outcome. Therefore, if the D+ estimate is very high, and at the same time the 1-N- estimate is very low, there will be a strong update towards the drug cue in this trial. But if the ratio between the two is similar, there will be a weaker update towards this drug relevant state. In this way, the overall update depends on the likelihood that the drug positive cue (D+) will predict the win, but also on the likelihood that the neutral negative cue (N-) will *not* predict the win (1-N-). Therefore, the larger the ratio/difference between the two likelihood parameters, the larger the update towards one relevant state over the other.

The probability estimate of a non-reversal (nR) between the two relevant states is another free parameter estimated for each participant. Here, nR represents the probability that no switch will occur, and 1-nR is the probably that a switch will occur. The greater the estimated probability of a non-reversal (nR), the more stable the participant believes the environment is, and the more they use their prior beliefs to make inferences. The transition probability between the two hidden states was encoded in a 2×2 matrix below:

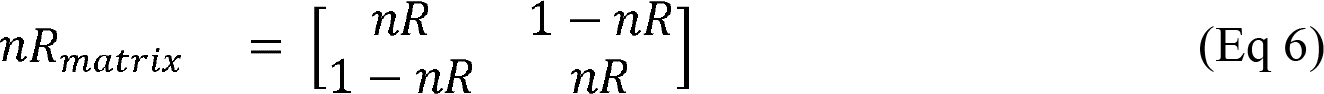

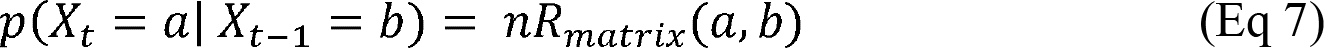

The nR could be between 0.5 and 1. A nR = 0.5 corresponds to the participant not using their prior beliefs at all, that is, the environment is estimated to have no stability/pattern. A nR = 1, corresponds to the participant only using their prior beliefs to make their inferences, with the environment being estimated as completely stable. This matrix is then multiplied by the prior belief of the relevant hidden state under both hypotheses (i.e., *p*(*X_t_* = 1 | *O*_1:*t*-1_) *and p*(*X*_*t*_ = 2 | *O*_1:*t*-1_)). Therefore, to weight the prior beliefs with estimated reversal probabilities, there is a multiplication of a 1×2 matrix (prior beliefs of the two hypotheses), with the 2×2 matrix (*nR_matrix_*). This multiplication allows for the prior beliefs to be weighted by the appropriate nR.

In sum, we estimated 5 free parameters per participant, and four of these are likelihood parameters, one for each cue. These indicated how much reliability each participant placed on each of the four cues (likelihood). The fifth free parameter is the non-reversal probability (nR). For details on the parameter recovery procedures, see *Supplementary Materials*. Importantly, we validated the parameter recovery procedure used here, through simulations using identical task condition/contingencies under the proposed model, with reliable recovery at the appropriate bounds for all 5 free parameters (see *Supplementary Materials*; Figure S1).

To determine internal representation updating based on the model estimated prior and posterior distributions, we calculated the Kullback-Leibler divergence (KLD). This KLD was calculated on a trial-by-trial basis and is the difference between the prior and posterior distribution (see equation 8 below).

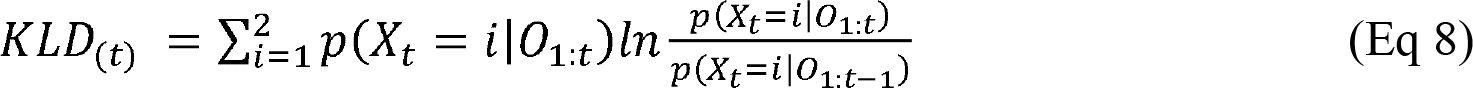

Based on this equation, if there is no difference between the prior and posterior distributions for that trial, the KLD will be 0, indicating no update to the internal representation. If the KLD is greater than 0, then there is an update of the internal representations. Importantly, the KLD computation gives the magnitude of the update, but does not give a direction of whether this update was towards the drug (H1; X = 1) or neutral (H2; X = 2) relevant state. However, the direction of each update can be determined based on the difference between the prior and posterior of the two hypotheses. For example, if the prior at trial t, for H1 and H2 is 0.8 and 0.2, respectively, and the posterior at trial t is 0.9 and 0.1, for H1 and H2, respectively, then the KLD calculated at trial t is updated towards the direction of H1, as the belief for H1 went from 0.8 to 0.9, and for H2 it went from 0.2 to 0.1. In this way, we treated KLD as a 2-dimentional vector, with magnitude and direction of the internal representation update.

### 2.9. Pupillometry pre-processing

The raw pupillometry data was preprocessed using a custom script adapted from Bogdanov (2021). We first identified blinks as values below 1. Next, we identified rapid/high speed deviations as well as any large gaps, which are variations usually caused due to the eye-tracker losing the eye momentarily or the participant moving out of range for the eye-tracker. All these blinks and large deviations were removed and replaced with a cubic interpolation (using the “interp1” function in MATLAB 2017b) for time sensitive analyses. We then baseline corrected (Carter & Luke, 2020; Mathôt et al., 2018; O’Reilly et al., 2019) the data, with the mean of 500ms prior to the epoch of interest. The “outcome” epoch was our main epoch of interest, and there was a blank screen prior to this (“anticipation” screen; see Figure 2). All pupil size data was z-scored per participant to allow between participant comparisons.

### 2.10. Analysis

All trials were split into two groups: 1) informative (where the two cues predicted opposite outcomes, allowing participants to infer which state they are likely in) and 2) uninformative trials (where both cues predicted the same outcome and did not allow participants to infer which state they are in). We further subdivided informative trials into four trial types; 1) *drug win* - where the win outcome was predicted by the drug cue, 2) *drug loss –* where the loss outcome was predicted by the drug cue, 3) *neutral win –* where the win outcome was predicted by the neutral cue, and finally, 4) *neutral loss –* where the loss outcome was predicted by the neutral cue. To test for significance, we used an ANOVA design with 3 factors, with 2 levels per factor. These included the factor of group (TUD and controls), cue type that predicted the outcome (drug or neutral) and outcome (win or loss).

The uninformative trials were subdivided into two trial types; 1) *win* – where both cues predicted the outcome win and, 2) *loss –* where both cues predicted the outcome loss. To test for significance for these uninformative trials, we used an ANOVA with factors of group (TUD and controls) and outcome (win or loss). In all analyses, t-tests were used to test for simple effects, adjusted for multiple comparisons using the Tukey method. Importantly, we did not include any “surprise” trials in these analyses. Lastly, the boxplot method was used to identify and exclude outliers, where participants with values +/- 3 times the interquartile range were excluded. All statistical analyses were done using RStudio 4.1.2. A complete list of results from the ANOVAs and t-tests are listed in the *Supplementary Materials*.

## 3. RESULTS

### 3.1. Model independent behavioral analysis

#### 3.1.1. Compared to the control group, people with a TUD misrepresented the drug and neutral cue-outcome relationships

Figure 3a shows the average rating towards the drug or neutral cue for informative trials, for each of the four trial types. Here, a rating of 1 indicates absolute confidence in the drug cue predicting the outcome, and a rating of 11 is absolute confidence in the neutral cue predicting the outcome. There was a significant main effect of cue type (F(1,176) = 1200.58, p = 2.2e-16), indicating that both groups did correctly attribute trials where drug cues predicted the outcome (“drug win” and “drug loss”) towards the drug cue, and vice versa for neutral predictive trials (“neutral win” and “neutral loss”). There was also a significant interaction between the cue type and group factors (F(1,176) = 19.20, p = 2.023e-05). Subsequent t-tests suggested that both groups equally attributed drug predicted wins to the drug cue (p = 0.27; “drug win” trials), however, when drug cues predicted a loss (“drug loss” trials), the TUD group rated more towards the neutral cue having predicted this outcome, compared to controls (p = 7e-04), as predicted by our SMMA theory. Further, and also in keeping with our SMMA prediction, when neutral cues predicted a win (“neutral win” trials), the TUD group rated more towards the drug cue having predicted this win outcome, compared to controls (p = 0.045). However, contrary to our prediction, when the neutral cue predicted a loss, the TUD group were also more likely to rate it towards the drug cue having predicted this loss (p = 0.03; “neutral loss”). Overall, compared to controls, the TUD group misrepresented the cue-outcome relationships in all trial types, except for when the drug cue predicted a win (“drug win” trials).

**Figure 3.**
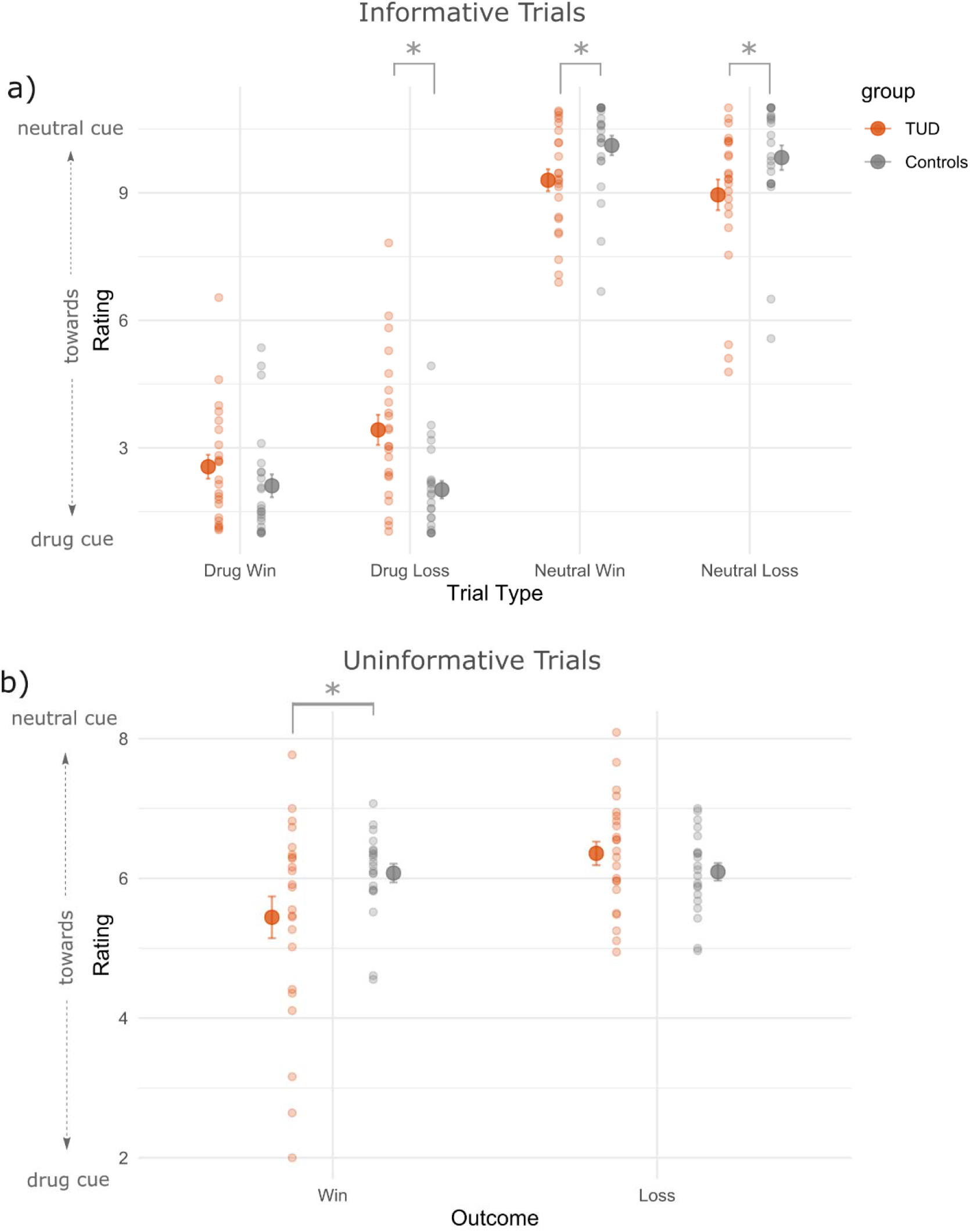
Ratings towards drug and neutral cues given the outcomes. Smaller dots are mean ratings per participant, with the larger dots being the overall mean +/- standard error of the mean. a) Ratings for informative trials. The TUD group performed similar to controls in attributing drug predictive wins to the drug cues (“drug win” trials) (p = 0.27). However, the TUD group attributed drug predictive losses more so towards the neutral cues, compared to the controls (“drug loss” trials; p = 0.0006). When neutral cues predicted a win, the TUD group was more likely to attribute this win to the drug cues (“neutral win” trials; p = 0.045). Lastly, however, when neutral cues predicted a loss, the TUD group was also more likely to attribute this towards the drug cues (“neutral loss” trials; p = 0.032). b) Ratings for uninformative trials. When neither cue predicted a win, the TUD group were more likely to attribute this win to the drug cues (p = 0.028). However, both groups performed similarly for the loss outcome (p = 0.35).

Figure 3b shows the average rating towards drug and neutral cues, but for uninformative trials, with “win” or a “loss” outcome. There was a significant main effect of outcome (F(1,84) = 5.48, p = 0.022) and a significant interaction between group and outcome ((F(1,84) = 5.03, p = 0.028). Subsequent t-tests revealed that compared to the control group, the TUD group was more likely to attribute uninformative wins to the drug cue (p = 0.028), consistent with our prediction, however, they did not perform differently to the control group for the loss outcome (p = 0.35).

#### 3.1.2. Compared to the control group, people with a TUD are biased in updating the drug and neutral cue-outcome relationships

Figure 4 shows the proportion of updates towards the drug cue. Updates for a given trial were calculated as the difference between the rating of the current trial (t), and the trial before (t-1). Therefore, to calculate proportion of updates towards the drug cue, we summed all the updates towards the drug cue, and divided this by the total updates (updates towards both drug and neutral cues). Figure 4a shows the proportion of updates towards the drug cue for informative trials. There was a significant main effect of cue type (F(1,176) = 1221.94, p = 2.2e-16), indicating that both groups did update more so towards the drug cue when they predicted the outcome, and towards the neutral cue when neutral cues predicted the outcome. There was also a significant group and cue type interaction (F(1,176) = 18.22, p = 3.218e-05). Subsequent t-tests suggested that both groups updated similarly for drug win (p = 0.28) and neutral win (p = 0.20) trials. However, when drug cues predicted a loss (“drug loss” trials), the TUD group updated less so towards the drug cue compared to the control group (p = 1e-04), in keeping with SMMA. However, inconsistent with SMMA, when neutral cues predicted a loss, the TUD group updated more towards the drug cue, compared to controls (p = 0.03). Overall, compared to controls, the TUD group had a bias in updating/learning the predictive relationships for the informative loss outcome, but not the informative win outcome. Figure 4b is the proportion of updates towards the drug cue for uninformative trials. There was a non-significant trend for the main effect of group (F(1,92 = 3.08, p = 0.08), with the trend being that the TUD group had a higher proportion of updates towards the drug cue for uninformative wins, but not for losses.

**Figure 4.**
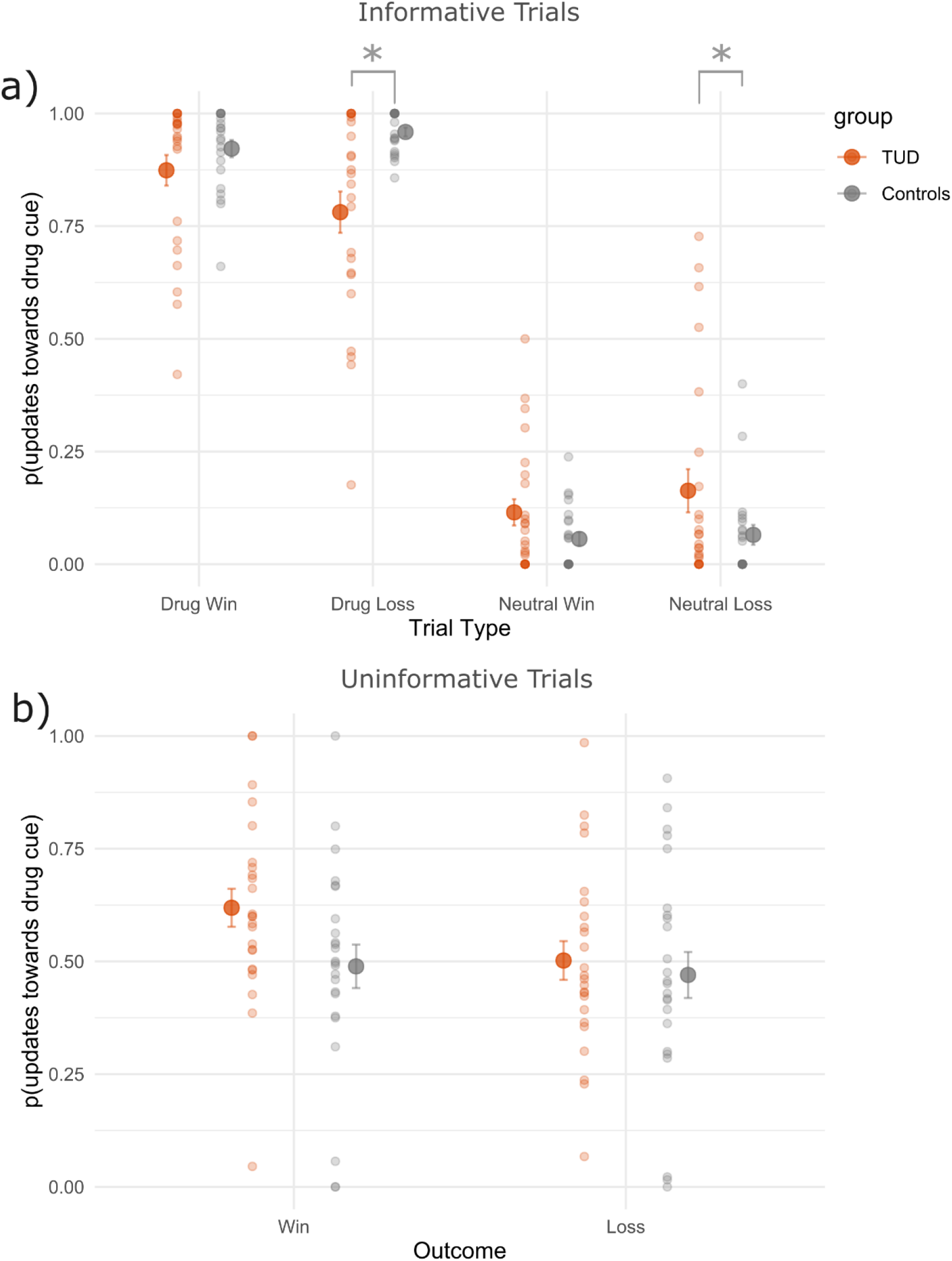
Proportion of updates towards the drug cue based on ratings. Smaller dots are mean updates per participant, with the larger dots being the overall mean +/- standard error of the mean. a) Proportion updates towards the drug cue for informative trials. Both groups updated similarly for the drug and neutral win trials. However, when losses were predicted by the drug cue, there were fewer updates towards the drug cue by the TUD group, compared to controls (“drug loss” trials; p = 1e-04). When neutral cues predicted the loss, the TUD group had more updates towards the drug cue, compared to controls (“neutral loss” trials; p = 0.030). b) Proportion updates towards the drug cue for uninformative trials. When neither cue predicted a win, the TUD group had a non-significant trend in updating towards the drug-cue.

### 3.2. Model based analyses

#### 3.2.1. Compared to the control group, people with a TUD generally had lower likelihood estimates

Figure 5 shows the recovered parameter estimates, which was optimised by minimising the difference between the observed and model generated rating behaviour (see *Supplementary Materials* for details). There was a significant main effect of group (F(1,42) = 9.34, p = 0.004), a significant main effect of parameter type (F(4,176) = 4.76, p = 0.001) and a significant interaction between group and parameter type (F(4,176) = 4.56, p = 0.001). Subsequent t-tests suggested that the TUD group had a lower likelihood for the win predictive drug cue (D+; p = 0.007), win predictive neutral cue (N+; p = 1e-4) and the loss predictive neutral cue (N-; 0.018) parameters. The loss predictive drug cue (D-) had similar likelihood for both groups (p = 0.41). Lastly, the non-reversal probability parameter (nR) was also similar for both groups with (mean +/- SE) of 0.76 +/- 0.027 for the TUD group and 0.80 +/- 0.027 for controls (not plotted; p = 0.28). Overall, the parameter estimates suggested that the TUD group generally had a low likelihood estimate for all cues, compared to the control group, except for drug predicting losses (D-).

**Figure 5.**
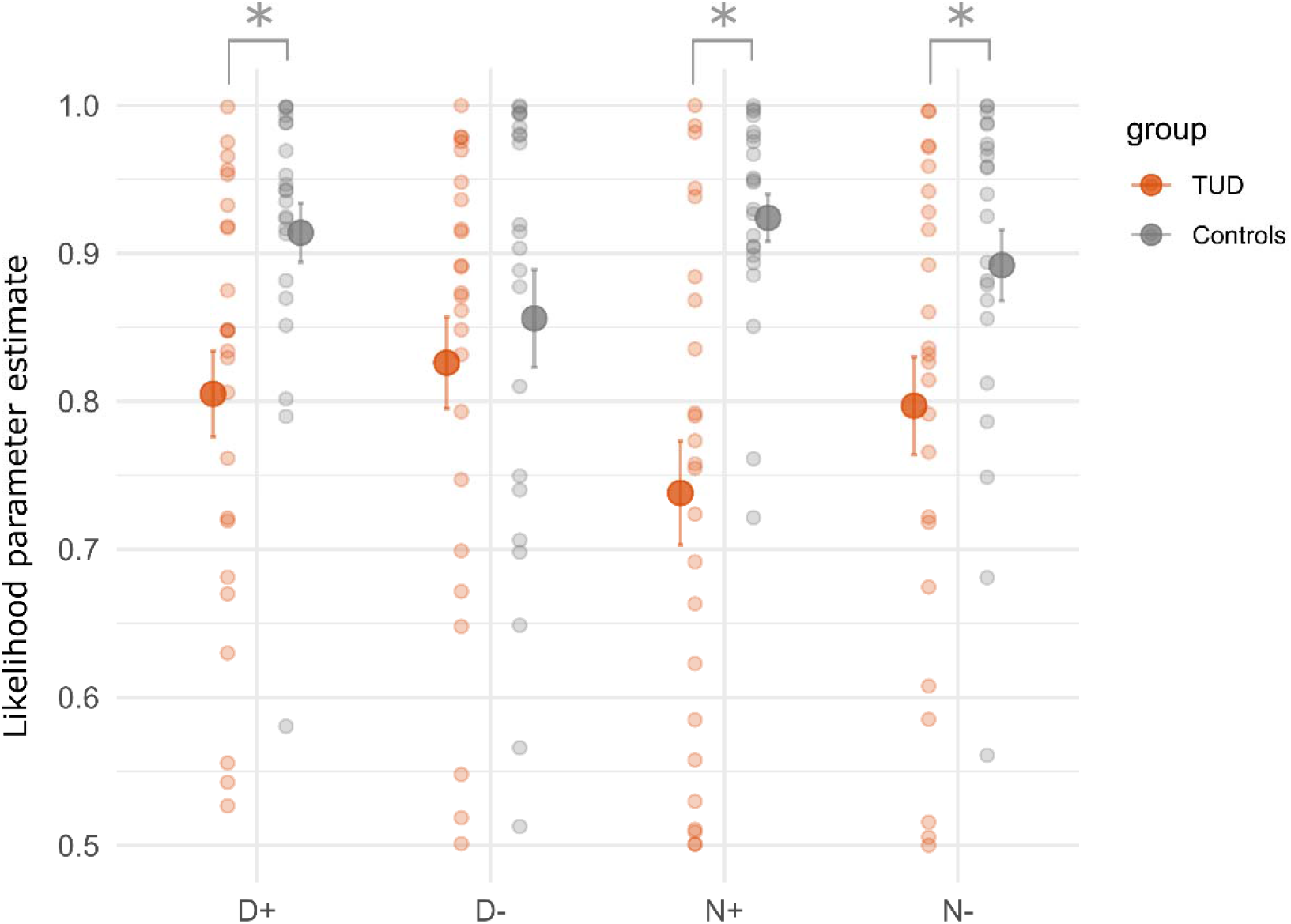
Recovered likelihood parameter estimates. Smaller dots parameter estimates per participant, with the larger dots being the overall mean +/- standard error of the mean. Compared to controls, the TUD group had a lower likelihood estimate for all parameters (p < 0.05), except for the D- parameter (p > 0.05). Abbreviations: D+ = drug predicting win likelihood, D- = drug predicting loss likelihood, N+ = neutral predicting win likelihood and N- = neutral predicting win likelihood.

#### 3.2.2. Differences in the likelihood parameter estimates between groups captured the bias in internal representation updating in people with a TUD

Here we used the recovered parameter estimates from each participant (see Figure 5) to estimate the prior and posterior distributions per participant on a trial-by-trial bases, using our Bayesian belief updating model. Based on these prior and posterior belief distributions, we calculated the trial-by-trial KLD magnitude and direction (update towards the drug or neutral relevant state; see *Computational Modelling* for details) for each participant. Figure 6a shows the total magnitude of updates (KLD) towards drug and neutral cues, for informative trials. In order to differentiate the direction of the update, we subtracted the total KLD towards drug cues with the total KLD towards neutral cues, for each participant. This subtraction made updates towards the drug cues positive, and updates towards the neutral cue, negative. We found a significant main effect of cue type (F(1,172) = 363, p = 2.2e-16), indicating that both groups updated more towards the drug cue when these were predictive of the outcome, and towards the neutral cue when they predicted the outcome. There was also a significant group and cue type interaction (F(1,172) = 21.19, p = 8.052e-06). Subsequent t-tests suggested that both groups updated their internal representations similarly for drug win (p = 0.06) and neutral win (p = 0.36) trials. When a loss was predicted by the drug cue, the TUD group had fewer updates towards the drug cue compared to controls (p = 0.001), in keeping with the SMMA theory. However, when a loss was predicted by the neutral cue, the TUD group updated more towards the drug cue, compared to the control group (p = 0.01), which is inconsistent with the SMMA theory. Overall, compared to the control group, the TUD group updated their internal representations less when drug and neutral cues predicted losses, but not wins.

**Figure 6.**
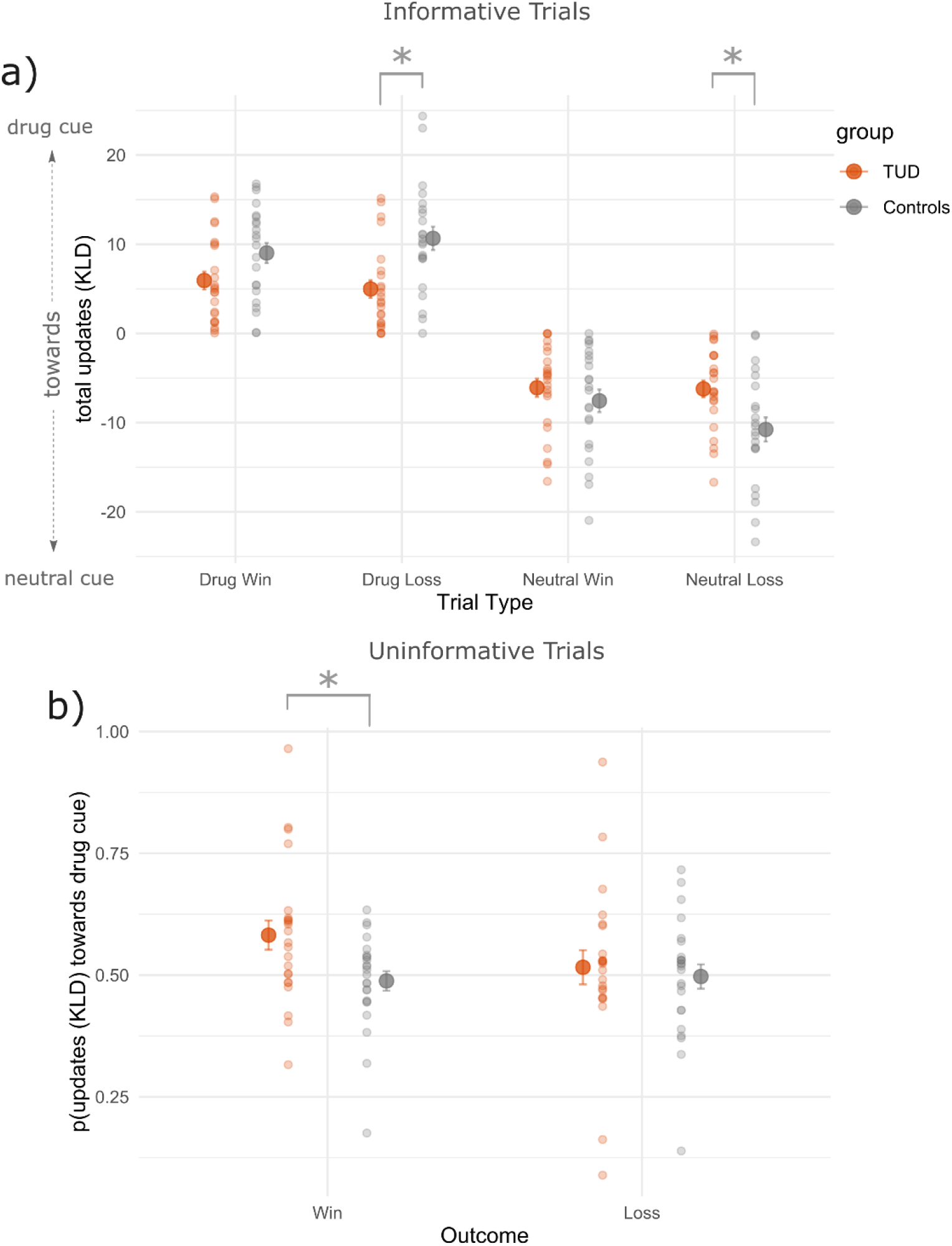
Internal representation updates using KLD. Smaller dots are mean updates per participant, with the larger dots being the overall mean +/- standard error of the mean. a) Total updates (KLD) towards the drug cue and neutral cues for informative trials. Positive values indicate that the update was towards the drug cue, and negative towards the neutral cue. Both groups updated similarly for the drug and neutral win trials. However, when losses were predicted by the drug cue the TUD group updated less towards the drug cue than controls (“drug loss” trials; p = 0.001). When neutral cues predicted the loss, the TUD group updated more towards the drug cue than controls (“neutral loss” trials; p = 0.01). b) Proportion updates towards the drug cue for uninformative trials. When neither cue predicted a win, the TUD group updated more towards the drug cue (p = 0.02). However, both groups performed similarly for the loss outcome (p = 0.63).

Figure 6b captures the proportion of internal representation updates (KLD) towards the drug cue for uninformative trials. This was calculated as the total KLD towards the drug cues divided by the total KLD/updates (towards both, drug and neutral cues). Based on the KLD proportions, there was a significant main effect of group (F(1,90) = 4.08, p = 0.046). Subsequent t-tests suggested that the TUD group updated more towards the drug cue during uninformative wins, when neither cue predicted the win outcome (p = 0.02). However, both groups updated similarly for uninformative loss outcomes (p = 0.63). Overall, the TUD group had a bias in updating their internal representations more towards the drug cue for wins, when neither cue predicted this win outcome.

In sum, these model-based results were consistent with the model independent behavioural results (see Figure 3), indicating that the differences in the parameter estimates between groups (Figure 5) contributed to the behavioural differences observed.

### 3.3. Eye-tracking analyses

#### 3.3.1. Gaze Behaviour

Figure 7a shows the proportion of gaze towards the drug cue for informative trials. This proportion was calculated as the time spent with the on the drug cue, divided by the total time (spent with the gaze on both drug and neutral cues). There was a significant three-way interaction between group, cue type, and outcome (F(1,114) = 5.68, p = 0.019). Subsequent t-tests suggested that the TUD group had a higher gaze proportion towards the drug cue for neutral loss trials (p = 0.03), compared to the control group. Both groups had a similar gaze proportion in the other informative trial types (p > 0.05). Figure 7b shows proportion of gaze towards the drug cue for uninformative trials. There was a non-significant trend for the main effect of group (F(1,78) = 3.61, p = 0.06).

**Figure 7.**
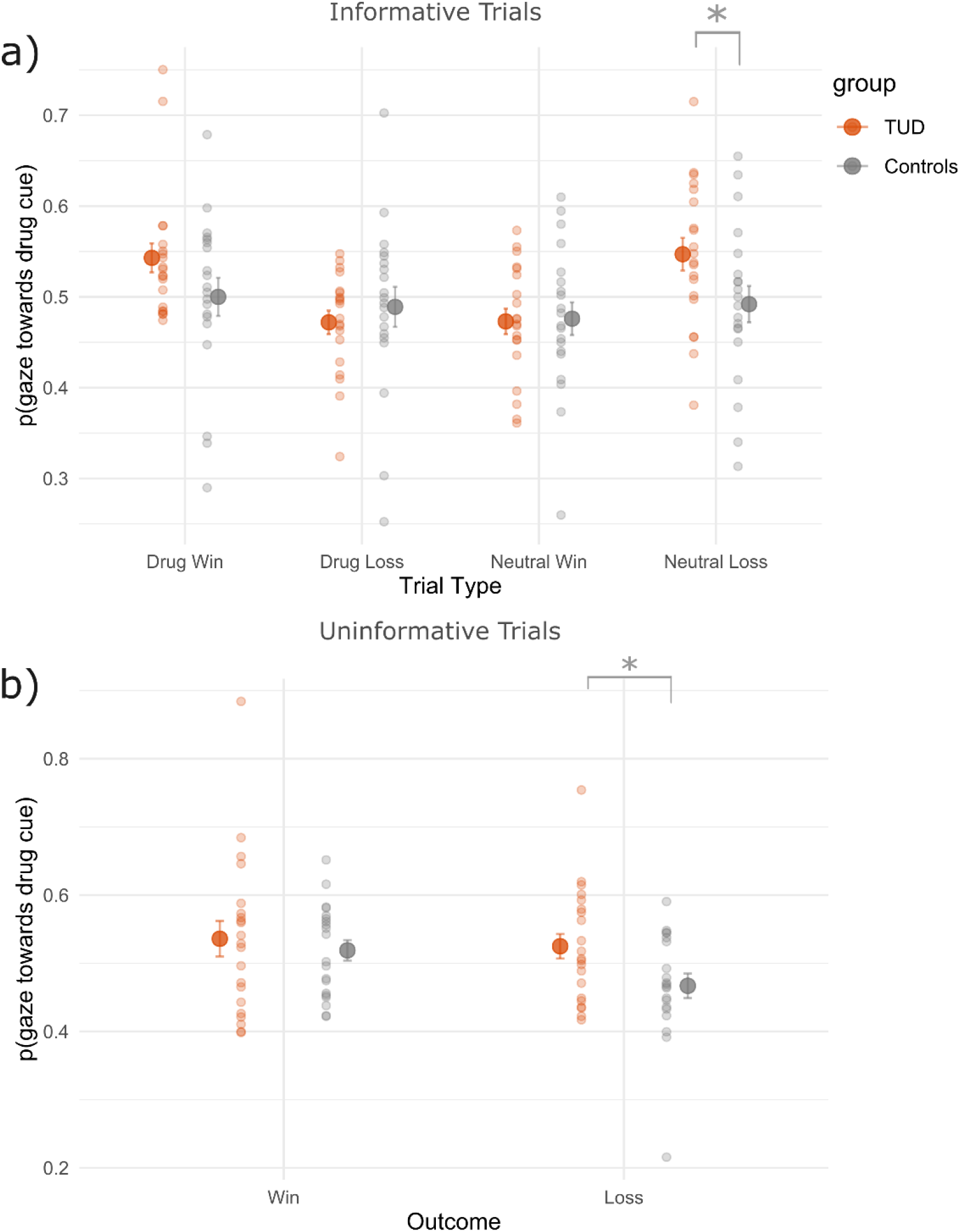
Gaze behaviour. Smaller dots are mean gaze proportions per participant, with the larger dots being the overall mean +/- standard error of the mean. a) Proportion of gaze towards the drug cue for informative trials. The TUD group had a higher gaze proportion towards the drug cue for the neutral loss trials (p = 0.03). There were no differences between groups for the other informative trials (p > 0.05). b) Proportion of gaze towards the drug cue for uninformative trials. Both groups had similar gaze proportions for win trials (p = 0.55). The TUD group had a higher gaze proportion towards the drug cue for loss trials, compared to controls (p = 0.04).

Next, we used a linear regression model to determine whether gaze proportion towards the drug cue had a linear relationship with internal representation updating (KLD) towards the drug cue. That is, does a longer gaze towards the drug cue predict larger KLD updating towards the drug cue? We performed a linear regression separately for each group and trial type and found a significant linear correlation for “neutral loss” informative trials, where the control group had a negative relationship between gaze towards the drug cue and KLD updates towards the drug cue (adjusted r^2^ = 0.21, p = 0.026; not plotted). This negative relationship suggested that for trials where the neutral cue predicted a loss, the more the gaze was towards the drug cue, the less they updated towards the drug cue. However, there were no other linear relationships between gaze proportions and internal representation updating, suggesting that eye gaze behaviour likely did not determine updates for most trial types in the current task, for both groups.

#### 3.3.2. In the TUD group (but not the control group), pupil size positively correlated with internal representation updating during informative drug win trials

Here, we used a linear regression model to correlate the mean pupil size at outcome phase with the mean updates (KLD) towards the drug cue. This linear regression was performed separately per group, for all trial types in the informative and non-informative trials. There were no significant correlations between pupil size and KLD updates towards the drug cue, except for the drug win informative trials (adjusted r^2^ = 0.44, p = 6.3e-04), for the TUD group (Figure 8a), but not for the control group (Figure 8b). When the drug cue predicted the win outcome, the TUD group showed larger pupil sizes for greater updates towards the drug cue.

**Figure 8.**
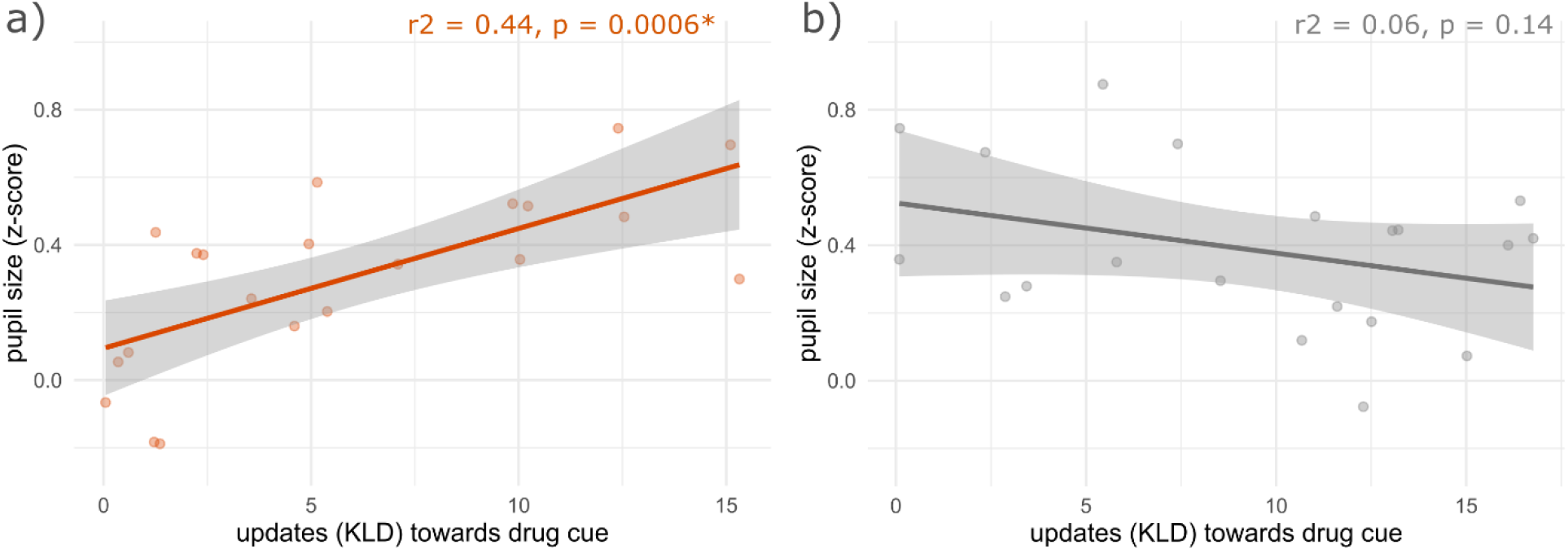
Pupil size positively correlated with updates (KLD) towards the drug cue for informative “drug win” trials for the a) TUD group, but not for b) the controls. Dots are means per participant, with the line +/ standard error estimated from a liner regression.

## 4. DISCUSSION

We investigated internal representation updating processes in people with a TUD and compared them to a non-smoking control group using a reversal learning task. The task had reversals in whether drug or neutral cues were relevant in predicting future outcomes (monetary wins or losses). Model independent and model-based behavioural results suggested that compared to the control group, the TUD group misrepresented drug and neutral cue-outcome relationships (Figure 3), with maladaptive internal representation updating processes (Figure 4 and 6). There was no linear relationship between internal representation updating (KLD), and eye behaviour (cue gaze and pupil size during the outcome epoch), with some exceptions (see Figures 7 and 8). Overall, we found key differences in how drug and neutral cue-outcome relationships are encoded and updated within the internal representation of people with a TUD and non-smoker controls.

### 4.1. Model independent behavioural results

Consistent with our hypotheses, compared to the control group, the TUD group misattributed drug predicted losses by rating more towards the neutral cue having predicted this loss outcome (Figure 3a). Further, when neutral cues predicted a win, the TUD group misattributed this win to the drug cue (Figure 3a). These results are consistent with our salience misattribution model (Kalhan et al., 2021), where people with a SUD are thought to misattribute drug cues as better predictors of positive outcomes, and neutral cues as better predictors of negative outcomes, even when both have equal predictive values. However, the TUD group also misattributed neutral predicted losses by rating more towards the drug cues, compared to the control group. Our SMMA theory predicted otherwise. We expected that compared to controls the TUD group would misattribute losses predicted by the neutral cue even more towards the neutral cues.

A key pattern across these rating results (Figure 3a) is that the TUD group generally performed worse than controls in all these informative trial types (except for “drug win” trials). Therefore, where the SMMA theory predicted the TUD group to have a bias in the direction of performing better than controls, the prediction was not met. An absence of a group difference specifically for these drug win trials suggested that the TUD group performed as well as controls only when the performance entailed forming a drug positive association, but not the other associations, which is consistent with our SMMA theory and other associative learning and memory theories of addiction (Di Chiara, 1999; Hyman, 2005; Keiflin & Janak, 2015; Redish, 2004; Torregrossa et al., 2011). However, these behavioural results also point towards a key limitation in our experimental design. Our task was relatively easy during informative trials, which may have led individuals in both groups to reach ceiling levels. This possibly diminished key behavioural differences between groups, especially in cases where the SMMA prediction of misattribution bias relied on the TUD group having better performance than controls.

When neither cue predicted the outcome (uninformative trials), the TUD group misattributed wins to drug cues (Figure 3b). This rating misattribution is consistent with our SMMA theory and suggested that there is a bias in the TUD group, where a pattern of drug cues predicting the positive outcome emerges, even when no such predictive pattern exists. However, contrary to our hypothesis, the TUD group did not misattribute uninformative losses more towards the neutral cue, compared to controls. In contrast to the informative trials, these uninformative trials are less susceptible to confounds based on participants reaching ceiling level performances as no predictive pattern exists here. Hence, these uninformative trials better reflect the biases between groups due to the salience misattribution processes, and less likely based on confounds due to the experimental design.

Model independent internal representation updating results suggested that the TUD group performed as well as controls in updating the informative win contingences but performed worse than controls in updating from the losses, for both drug and neutral loss trials (Figure 4a). Specifically, when the drug cue predicted a loss outcome, the TUD group updated less towards the drug cue having predicted the loss, compared to the control group. The reduced updating towards the drug loss cue is consistent with the SMMA theory, which predicted a misaligned internal representation generated with reduced updating/learning towards the drug cue being predictive of negative outcomes. However, contrary to what we had expected, when the neutral cue predicted the loss (“neutral loss” trials), the TUD group updated more towards the drug cue having predicted the loss. Therefore, the internal representation is also misaligned in attributing neutral predicted losses towards the drug cue, which is inconsistent with the SMMA theory.

During these neutral loss trials, the neutral negative cue is presented next to the drug positive cue. Therefore, one interpretation for updating more towards the drug positive cue here, may be a compulsive-like habitual responding towards the drug positive cue, irrespective of the outcome. This interpretation is consistent with dual-processing theories of addiction (Dalley et al., 2011; Everitt & Robbins, 2005, 2013, 2016; Lüscher et al., 2020; Redish et al., 2008) which suggests that there may be some instances when the TUD group seeing the drug positive cue released a situation à action habitual chain (see Redish et al., (2008) Vulnerability 7), that leads to the action of rating towards the drug cue, irrespective of the outcome. Alternately, because the reduced internal representation updating is specific to informative loss outcomes, the TUD group may generally be worse at learning the loss contingencies, which is consistent with several other empirical other accounts (Carey et al., 2015; Duehlmeyer et al., 2018; Duehlmeyer & Hester, 2019; Forman et al., 2004; Franken et al., 2007; Hester et al., 2007, 2009, 2012; Hester, 2012). Our findings add to this body of work in that this impaired error learning is not dependent on the cue type (drug or neutral) and may be a general phenomenon within the SUD population.

During uninformative trials, when neither cue predicted a win, the TUD group attributed wins more to the drug cue (Figure 3-4b). The TUD group may be more susceptible to forming falsely strong drug-win associations during periods where there are no obvious predictive patterns but may be able to overcome this bias during the informative trials, where predictive patterns do exist. Unlike the informative loss trials, there was no bias in updating from uninformative losses. In sum, the findings for uninformative trials support a salience misattribution effect consistent with the SMMA theory, with TUD misattributing relevance of drug cues in predicting positive outcomes. However, we did not find evidence for a misattribution of neutral cues predicting negative outcomes.

Overall, the TUD group showed misattribution of wins and losses. The TUD group was generally worse than controls at attributing credit to the correct cue in all informative trial types except for when the drug cues predicted a win – possibly indicating that the drug-win association was the easiest to learn for the TUD group, consistent with our SMMA theory. Model independent updating results suggested that the TUD group updated similarly to controls for informative win trials, but not during the informative loss trials, with fewer updates towards the correct cue type for both drug and neutral loss trials. These results indicated that several decision-making systems may be involved, in addition to salience misattribution effects, including compulsive-like responding towards the drug positive cue, as well as a general aberration in learning from negative outcomes, irrespective of the cue type. During uninformative trials, the TUD group revealed a bias where they misattributed and updated more towards the drug cue as predictors of the wins, but there was no updating bias for uninformative losses. Therefore, a key finding, consistent with the SMMA theory, was that in the absence of a predictive pattern, the TUD group is susceptible to falsely forming a drug-win association. In sum, these model independent results suggested key behavioural differences between groups, but also possible task limitations. However, to gain a better understating of the underlying latent processes influencing the behavioural results, we utilized computational modelling using our Bayesian belief updating model (see below).

### 4.2. Model-based behavioural results

The recovered parameter estimates suggested that the TUD group had a lower likelihood estimate for all cue types, except for the drug negative cue (D-), compared to controls (Figure 5). We calculated the KLD on a trial-by-trial basis for every participant. The KLD updating results were consistent with the model independent updating results, where the TUD group updated less towards the correct cue during informative loss trials but had similar performance to controls for informative win trials, irrespective of the cue type (Figure 6a). Also consistent with model independent updating results, KLD updating for the TUD group was greater towards the drug cue for uninformative drug wins, but there were no group differences in updating during uninformative losses (Figure 6b). Given the consistency between the model-based and independent results, it is likely that the recovered likelihood parameter estimates are contributing to the behavioural differences observed between groups.

Most likelihood parameter estimates were lower for the TUD group, compared to the control group, explaining reduced updating in general by the TUD group during informative trials. The true overall likelihood estimate in the task for all cues were 0.8. Therefore, it may seem paradoxical that the TUD group had a closer likelihood estimate to the true task than the control group, yet the TUD group still had misaligned internal representation updating. However, this can simply be explained by the surprise trials. Given that reversals occurred very quickly in our task (every 8-12 trials), with half of these trials being uninformative trials, it was difficult for participants to distinguish between incongruent surprise trials and informative non-surprise trials (as they were both incongruent trials, where the two cues predicted opposite outcomes). Therefore, both groups treated incongruent surprise trials just as if they were informative trials and accordingly updated their internal representations (see Figure S2, supplementary materials). These updates during surprise trials caused both groups to have higher likelihood estimates. However, the control group correctly updated more than the TUD group during the informative trials (Figure 4a and 6a), indicating that they did form more accurate internal representations.

A key finding was that the TUD group had an especially low likelihood estimate of the neutral positive cue predicting a positive outcome (N+). This low N+ value also meant that these neutral positive cues had a relatively high likelihood estimate of, incorrectly, predicting a negative outcome (1-N+). Therefore, the TUD group updated less towards the drug negative cue during drug loss trials, not because they had a low likelihood estimates for the drug negative cue (D-), but because they had a relatively high likelihood estimate of the neutral positive cue in also predicting the loss outcome (1-N+). Consequently, as the SMMA theory predicted, the internal representation of the TUD group is misaligned towards the drug cues being less predictive of the loss. Modelling results added that the low N+ estimate by the TUD group was a key latent variable in producing this misaligned internal representation. The TUD group’s low N+ value estimate can also explain the bias found during uninformative win trials, where the TUD group updated more towards the drug positive cue in predicting the uninformative wins. The TUD group had a lower likelihood estimate of the neutral positive cue predicting a win (N+) compared to the likelihood estimate of the drug positive cue in predicting the win (D+). As a result, updating was biased towards the drug positive cue having predicted this uninformative win. Consequently, as the SMMA theory predicted, the internal representation of the TUD group was misaligned towards drug cues being better predictor of wins, even when no predictive patterns existed.

In sum, computational modelling suggested that the low N+ likelihood estimate of the TUD group was a key latent variable contributing to the behavioural results. Due to this low N+ value, we see a salience misattribution effect where 1) there are fewer updates by the TUD group towards the drug negative cue when they predict a loss during informative drug loss trials, and 2) there are more updates towards the drug positive cue predicting the win outcome, during uninformative win trials. Collectively, these findings support a key prediction of the SMMA theory where maladaptive updating produces a misaligned internal representation in people with a SUD such that non-drug cues are worse predictors of wins, but better predictors of losses. The critical factor explaining these results was that the neutral positive cue for the TUD group had a very low likelihood estimate in predicting the win (N+), and consequently, a relatively higher likelihood estimate in predicting a loss (1-N+).

### 4.3. Eye-tracking results

The gaze behaviour results suggested that the TUD group, compared to controls, had a higher gaze proportion towards the drug cue during neutral loss informative trials, and uninformative loss trials (Figure 7). These results suggest that the TUD group may have had greater visual attention towards the drug cue, before losses were observed. However, these gaze proportions did not correlate with KLD internal representation updating. Therefore, internal representation updating processes in the present task are unlikely related to visual salience attentional processes (Lambert et al., 2018), but more likely cognitive/motivational-based salience attentional processes.

Our SMMA theory predicts the involvement of these anterior cingulate cortex and dopaminergic processes in assigning a greater salience to cues with higher estimated predictive values and producing greater updating from these as a result (Akaishi et al., 2016; Kolling, Behrens, et al., 2016; Kolling, Wittmann, et al., 2016; Nour et al., 2018; O’Reilly et al., 2013; Schwartenbeck et al., 2016). We speculate that these salience-related neural processes may be involved in the current task, possibly explaining the misaligned internal representation updating in people with a TUD. However, neuroimaging data is needed to support or oppose our speculation.

There was no relationship between pupil size and KLD updates, except for the TUD group during informative drug win trials. During these trials where the drug cue predicted a win, the larger the pupil size during the win outcome epoch, the greater the KLD updating by the TUD group (Figure 8a). Such a linear relationship between pupil size and KLD updates was not found in the control group (Figure 8b). Pupil size generally correlates with greater attentional processing (Gabay et al., 2011), and has also correlated with KLD updating a similar task involving multiple cues (Hämmerer et al., 2019). Further, Zénon (2019) proposed that the many seemingly disparate decision-making related associations (e.g., uncertainly, learning rate, volatility) with pupil size may be explained by pupil size being generally associated with information gain (i.e., the KLD). Consistent with these accounts, we find that the TUD group’s pupil size positively correlated with KLD. However, this was only for drug predicted wins in the TUD group but not for other trials, nor for the control group. One possible explanation is that our task being very passive, and relatively easy, did not elicit large enough variations in pupil size, which may have hinder further relationships with KLD. It is possible that in the TUD group drug predicted wins may have captured more attentional resources/variability.

### 4.4. Implications

A key finding was that the TUD group had a very low likelihood estimate for neutral cues in predicting the positive outcome (N+). The low N+ estimate here led to the misaligned internal representation updating processes in the TUD group where losses predicted by the drug cue produced fewer updates towards forming the drug-loss predictive association. Further, the low N+ estimate also led to more updates towards forming the drug-win predictive association during uninformative win trials. An implication of misaligned updating is likely the formation of a maladaptive internal representation, where drug cues are more predictive of positives, but less predictive of negative outcomes. Therefore, based on this maladaptive internal representation, it is more likely for the individual to further engage in drug taking actions, as they are misattributed as better predictors of positive outcomes.

Our finding of the reduced N+ value for the TUD group is also consistent with previous accounts suggesting that people with a SUD usually find non-drug positive cues less engaging/salient, including those associated with primary rewards (Garavan et al., 2000; Verdejo-Garcia et al., 2018; Volkow et al., 1997; Volkow & Li, 2004). However, our finding goes further to suggest that the reduced salience towards the non-drug positive cues may be because these cues have a reduced estimated likelihood of predicting positive outcomes, which we have captured through our Bayesian belief updating model. Consequently, treatments such as cognitive behaviour therapy where associations between non-drug cues and positive outcomes are strengthened may help restore the misaligned internal representation, and possibly aid in reducing drug related behaviours. The current task, in combination with other similar tasks and diagnostic tools/questionnaires may also be used to identify those with such maladaptive associations, with treatments more personalised based on restoring such associations. In sum, we used strong theoretical hypotheses in concert with computational modelling and eye-tracking to gain a better understanding of the internal representation updating processes of people with a TUD.

## Notes

### Competing Interest Statement

The authors have declared no competing interest.

