## Supplementary Materials for "People with a tobacco use disorder misattribute non-drug cues as worse predictors of positive outcomes compared to drug cues"

*Parameter recovery*

We estimated parameters in two steps using two separate functions within MATLAB2017b. In all these steps, we estimated parameters that minimized the sum squared of the difference between true and model generated behavior (i.e., minimized the difference between model generated and true behavior, and thereby maximized the correlation between model estimated and true behavior). All parameters were estimated separately for each participant. The first step was using a genetic algorithm using the “ga” function. This generic algorithm gave a fast, but crude parameter estimate. Therefore, we used the parameter estimates from this genetic algorithm as the starting point for the next step which was using the “fmincon” function, which fine-tuned the parameter estimates. We repeated this procedure 10 times per participant, and reported the parameter estimates that gave the lowest difference between true and model generated behavior. Importantly, we were able to validate this parameter recovery procedure, and justify the bounds placed, through simulations that generated behavior using the model, using the same task contingencies as the true task (see Figure S1). Overall, we showed reliable recovery of all four likelihood parameters between 0.5 and 1, and between 0.6 and 1 for the non-reversal probability parameter. These were the bounds placed for the “ga” and “fmincon” functions when recovering parameters.

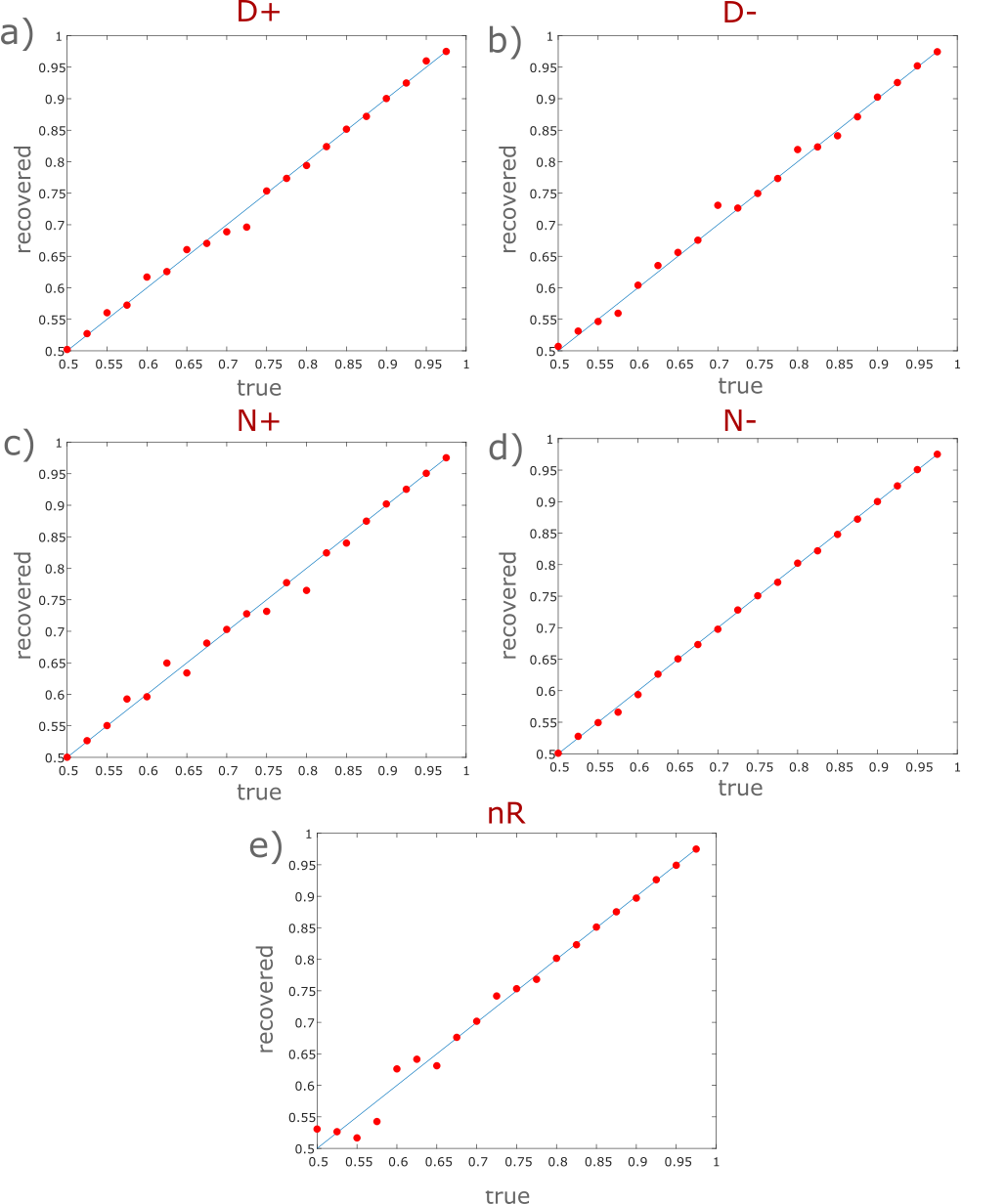

***Figure S1.*** *True and recovered values from each of the 5 parameters estimates*. *Abbreviations: D+ = drug positive likelihood, D- = drug negative likelihood, N+ = neutral positive likelihood, N- = neutral negative likelihood, nR = non-reversal probability.*

*Supplementary Results*

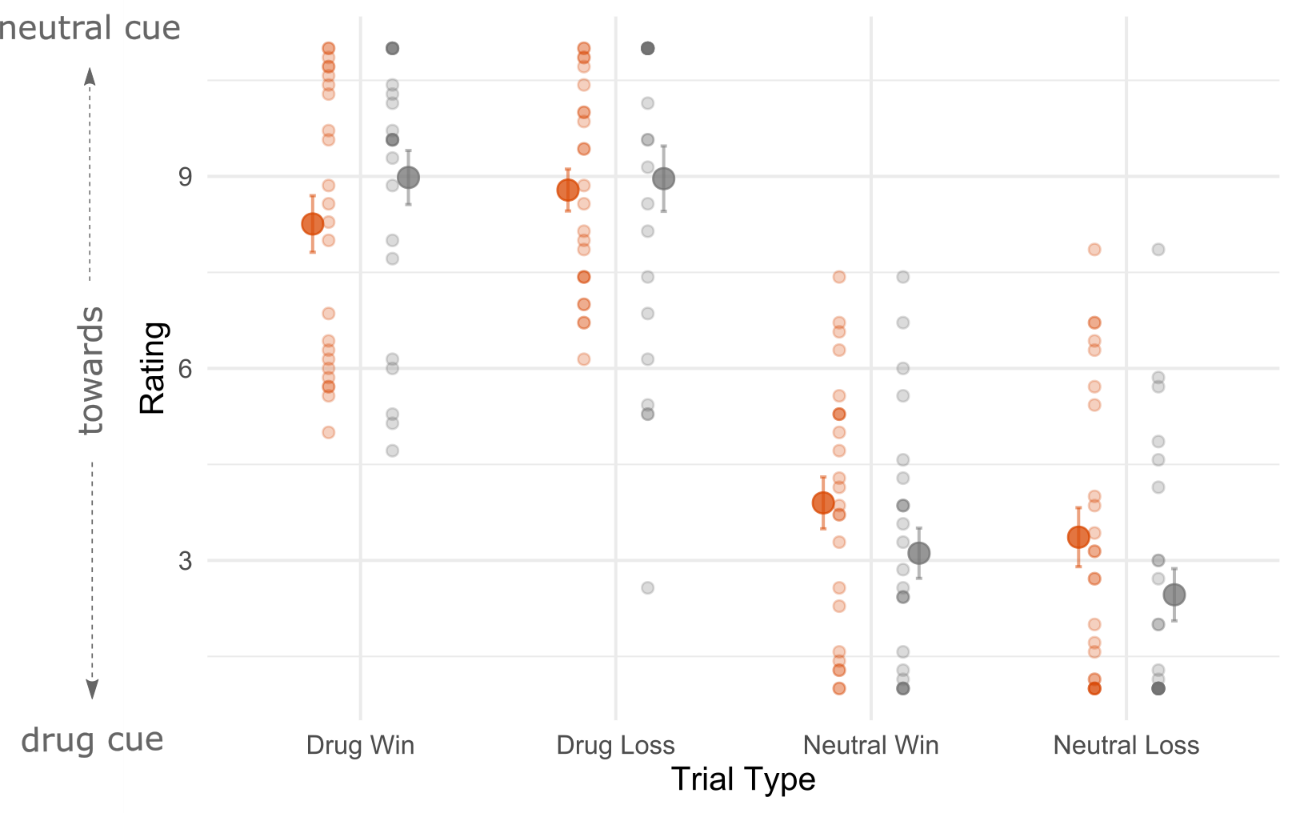

***Figure S2.*** *Rating scores for surprise informative trials.* During surprise trials, both groups rated towards the opposite cue having predicted the outcome (main effect of cue type; F(1,184) = 341.08, p = 2e-16). This main effect of cue type indicated that both groups updated from surprise trials, just as if they were informative trials.

*Analysis – AVOVA and t-test tables*

```
Key:
Group- 1 = smokers, 2 = controls
Outcome - 1 = win, 2 = loss
Cue type - 1 = drugs, 2 = neutral
ptype (parameter type) - 1 = drug positive, 2 = drug negative,
3 = neutral positive, 4 = neutral negative


Figure 3a - informative ratings
|                       |       Sum Sq|      Mean Sq| NumDF| DenDF|      F value|    Pr(>F)|
|:----------------------|------------:|------------:|-----:|-----:|------------:|---------:|
|group                  |    0.0735429|    0.0735429|     1|   176|    0.0389344| 0.8438063|
|outcome                |    0.0599559|    0.0599559|     1|   176|    0.0317413| 0.8588015|
|cue_type               | 2267.7709697| 2267.7709697|     1|   176| 1200.5834738| 0.0000000|
|group:outcome          |    2.3239311|    2.3239311|     1|   176|    1.2303153| 0.2688590|
|group:cue_type         |   36.2575491|   36.2575491|     1|   176|   19.1951545| 0.0000202|
|outcome:cue_type       |    5.6775801|    5.6775801|     1|   176|    3.0057748| 0.0847195|
|group:outcome:cue_type |    2.9740531|    2.9740531|     1|   176|    1.5744972| 0.2112188|

|contrast        |outcome |cue_type |   estimate|        SE|  df|   t.ratio|   p.value|
|:---------------|:-------|:--------|----------:|---------:|---:|---------:|---------:|
|group1 - group2 |1       |1        |  0.4487578| 0.4052794| 176|  1.107280| 0.2696839|
|group1 - group2 |2       |1        |  1.4068323| 0.4052794| 176|  3.471265| 0.0006517|
|group1 - group2 |1       |2        | -0.8183230| 0.4052794| 176| -2.019158| 0.0449882|
|group1 - group2 |2       |2        | -0.8773292| 0.4052794| 176| -2.164752| 0.0317535|


Figure 3b – non-informative ratings
|              |    Sum Sq|   Mean Sq| NumDF| DenDF|  F value|    Pr(>F)|
|:-------------|---------:|---------:|-----:|-----:|--------:|---------:|
|group         | 0.7497794| 0.7497794|     1|    84| 0.854448| 0.3579449|
|outcome       | 4.7888667| 4.7888667|     1|    84| 5.457388| 0.0218685|
|group:outcome | 4.4129101| 4.4129101|     1|    84| 5.028949| 0.0275610|

|contrast        |outcome |   estimate|        SE| df|    t.ratio|   p.value|
|:---------------|:-------|----------:|---------:|--:|----------:|---------:|
|group1 - group2 |1       | -0.6331337| 0.2827331| 84| -2.2393331| 0.0277768|
|group1 - group2 |2       |  0.2635315| 0.2827331| 84|  0.9320856| 0.3539636|


Figure 4a – informative updates (model independent)
|                       |     Sum Sq|    Mean Sq| NumDF| DenDF|      F value|    Pr(>F)|
|:----------------------|----------:|----------:|-----:|-----:|------------:|---------:|
|group                  |  0.0138890|  0.0138890|     1|   176|    0.6015931| 0.4390118|
|outcome                |  0.0000041|  0.0000041|     1|   176|    0.0001770| 0.9894006|
|cue_type               | 28.2110363| 28.2110363|     1|   176| 1221.9420700| 0.0000000|
|group:outcome          |  0.0231302|  0.0231302|     1|   176|    1.0018699| 0.3182326|
|group:cue_type         |  0.4205442|  0.4205442|     1|   176|   18.2155914| 0.0000322|
|outcome:cue_type       |  0.0367801|  0.0367801|     1|   176|    1.5931067| 0.2085528|
|group:outcome:cue_type |  0.0829395|  0.0829395|     1|   176|    3.5924672| 0.0596812|


|contrast        |outcome |cue_type |   estimate|        SE|  df|   t.ratio|   p.value|
|:---------------|:-------|:--------|----------:|---------:|---:|---------:|---------:|
|group1 - group2 |1       |1        | -0.0481510| 0.0448483| 176| -1.073641| 0.2844534|
|group1 - group2 |2       |1        | -0.1780459| 0.0448483| 176| -3.969956| 0.0001046|
|group1 - group2 |1       |2        |  0.0582558| 0.0448483| 176|  1.298951| 0.1956598|
|group1 - group2 |2       |2        |  0.0983702| 0.0448483| 176|  2.193397| 0.0295896|


Figure 4b - non-informative updates (model independent)
|              |    Sum Sq|   Mean Sq| NumDF| DenDF|  F value|    Pr(>F)|
|:-------------|---------:|---------:|-----:|-----:|--------:|---------:|
|group         | 0.1577982| 0.1577982|     1|    92| 3.084827| 0.0823533|
|outcome       | 0.1102344| 0.1102344|     1|    92| 2.154993| 0.1455178|
|group:outcome | 0.0583335| 0.0583335|     1|    92| 1.140373| 0.2883677|

|contrast        |outcome |  estimate|        SE| df|   t.ratio|   p.value|
|:---------------|:-------|---------:|---------:|--:|---------:|---------:|
|group1 - group2 |1       | 0.1303866| 0.0652897| 92| 1.9970461| 0.0487749|
|group1 - group2 |2       | 0.0317852| 0.0652897| 92| 0.4868325| 0.6275349|


Figure 5 - recovered parameters
|            |    Sum Sq|   Mean Sq| NumDF|     DenDF|  F value|    Pr(>F)|
|:-----------|---------:|---------:|-----:|---------:|--------:|---------:|
|group       | 0.0910590| 0.0910590|     1|  42.83618| 9.347371| 0.0038384|
|ptype       | 0.1854961| 0.0463740|     4| 176.88352| 4.760379| 0.0011351|
|group:ptype | 0.1776535| 0.0444134|     4| 176.88352| 4.559113| 0.0015776|

|contrast        |ptype |   estimate|        SE|       df|    t.ratio|   p.value|
|:---------------|:-----|----------:|---------:|--------:|----------:|---------:|
|group1 - group2 |1     | -0.1117500| 0.0406024| 110.9999| -2.7523019| 0.0069142|
|group1 - group2 |2     | -0.0331815| 0.0406024| 110.9999| -0.8172317| 0.4155467|
|group1 - group2 |3     | -0.1888844| 0.0406024| 110.9999| -4.6520526| 0.0000091|
|group1 - group2 |4     | -0.0977144| 0.0406024| 110.9999| -2.4066171| 0.0177516|
|group1 - group2 |5     | -0.0444643| 0.0406024| 110.9999| -1.0951163| 0.2758362|


Figure 6a - informative updates (model-based)
|                       |       Sum Sq|      Mean Sq| NumDF| DenDF|     F value|    Pr(>F)|
|:----------------------|------------:|------------:|-----:|-----:|-----------:|---------:|
|group                  | 2.127799e+01| 2.127799e+01|     1|   172|   0.7342062| 0.3927154|
|outcome                | 1.994252e+01| 1.994252e+01|     1|   172|   0.6881252| 0.4079526|
|cue_type               | 1.053417e+04| 1.053417e+04|     1|   172| 363.4861157| 0.0000000|
|group:outcome          | 6.814888e-01| 6.814888e-01|     1|   172|   0.0235151| 0.8783050|
|group:cue_type         | 6.141092e+02| 6.141092e+02|     1|   172|  21.1900984| 0.0000081|
|outcome:cue_type       | 4.558381e+01| 4.558381e+01|     1|   172|   1.5728886| 0.2114898|
|group:outcome:cue_type | 9.025125e+01| 9.025125e+01|     1|   172|   3.1141578| 0.0793895|

|contrast        |outcome |cue_type |  estimate|       SE|  df|    t.ratio|   p.value|
|:---------------|:-------|:--------|---------:|--------:|---:|----------:|---------:|
|group1 - group2 |1       |1        | -3.089442| 1.605415| 172| -1.9243887| 0.0559565|
|group1 - group2 |2       |1        | -5.676330| 1.605415| 172| -3.5357401| 0.0005226|
|group1 - group2 |1       |2        |  1.467645| 1.605415| 172|  0.9141841| 0.3618999|
|group1 - group2 |2       |2        |  4.546901| 1.605415| 172|  2.8322280| 0.0051745|


Figure 6b – non-informative updates (model-based)
|              |    Sum Sq|   Mean Sq| NumDF| DenDF|  F value|    Pr(>F)|
|:-------------|---------:|---------:|-----:|-----:|--------:|---------:|
|group         | 0.0757166| 0.0757166|     1|    90| 4.075303| 0.0464906|
|outcome       | 0.0188757| 0.0188757|     1|    90| 1.015948| 0.3161844|
|group:outcome | 0.0326269| 0.0326269|     1|    90| 1.756081| 0.1884685|

|contrast        |outcome |  estimate|        SE| df|   t.ratio|   p.value|
|:---------------|:-------|---------:|---------:|--:|---------:|---------:|
|group1 - group2 |1       | 0.0940447| 0.0397736| 90| 2.3645014| 0.0202039|
|group1 - group2 |2       | 0.0195060| 0.0397736| 90| 0.4904251| 0.6250267|


Figure 7a - informative gaze
|                       |    Sum Sq|   Mean Sq| NumDF| DenDF|    F value|    Pr(>F)|
|:----------------------|---------:|---------:|-----:|-----:|----------:|---------:|
|group                  | 0.0119867| 0.0119867|     1|    38|  1.9736285| 0.1681855|
|outcome                | 0.0001647| 0.0001647|     1|   114|  0.0271129| 0.8695031|
|cue_type               | 0.0006200| 0.0006200|     1|   114|  0.1020819| 0.7499302|
|group:outcome          | 0.0000076| 0.0000076|     1|   114|  0.0012548| 0.9718044|
|group:cue_type         | 0.0015816| 0.0015816|     1|   114|  0.2604140| 0.6108225|
|outcome:cue_type       | 0.0733624| 0.0733624|     1|   114| 12.0792665| 0.0007221|
|group:outcome:cue_type | 0.0344691| 0.0344691|     1|   114|  5.6754085| 0.0188593|

|contrast        |outcome |cue_type |   estimate|        SE|       df|    t.ratio|   p.value|
|:---------------|:-------|:--------|----------:|---------:|--------:|----------:|---------:|
|group1 - group2 |1       |1        |  0.0426146| 0.0253093| 150.7834|  1.6837513| 0.0942989|
|group1 - group2 |2       |1        | -0.0169688| 0.0253093| 150.7834| -0.6704568| 0.5035922|
|group1 - group2 |1       |2        | -0.0035196| 0.0253093| 150.7834| -0.1390652| 0.8895842|
|group1 - group2 |2       |2        |  0.0543178| 0.0253093| 150.7834|  2.1461585| 0.0334604|


Figure 7b - non-informative gaze
|              |    Sum Sq|   Mean Sq| NumDF| DenDF|  F value|    Pr(>F)|
|:-------------|---------:|---------:|-----:|-----:|--------:|---------:|
|group         | 0.0289756| 0.0289756|     1|    78| 3.607984| 0.0611996|
|outcome       | 0.0200781| 0.0200781|     1|    78| 2.500083| 0.1178871|
|group:outcome | 0.0087043| 0.0087043|     1|    78| 1.083841| 0.3010562|

|contrast        |outcome |  estimate|        SE| df|   t.ratio|   p.value|
|:---------------|:-------|---------:|---------:|--:|---------:|---------:|
|group1 - group2 |1       | 0.0169950| 0.0279996| 78| 0.6069752| 0.5456302|
|group1 - group2 |2       | 0.0582189| 0.0279996| 78| 2.0792800| 0.0408778|


Figure 8a - smokers pupil linear regression (drug win trials)
Residuals:
    Min      1Q  Median      3Q     Max
-6.4501 -2.3117  0.6271  1.6833  9.4299

Coefficients:
             Estimate Std. Error t value Pr(>|t|)
(Intercept)     1.946      1.286   1.514 0.146603
sm_DW_p$mean   13.179      3.230   4.080 0.000638 ***
---
Signif. codes:  0 ‘***’ 0.001 ‘**’ 0.01 ‘*’ 0.05 ‘.’ 0.1 ‘ ’ 1

Residual standard error: 3.76 on 19 degrees of freedom
Multiple R-squared:  0.467,	Adjusted R-squared:  0.4389
F-statistic: 16.65 on 1 and 19 DF,  p-value: 0.0006382


Figure 8b - controls pupil linear regression (neutral win trials)
Residuals:
    Min      1Q  Median      3Q     Max
-9.3780 -3.9597  0.2911  3.5670  8.2960

Coefficients:
             Estimate Std. Error t value Pr(>|t|)
(Intercept)    12.246      2.297   5.332 4.55e-05 ***
co_DW_p$mean   -7.764      5.077  -1.529    0.144
---
Signif. codes:  0 ‘***’ 0.001 ‘**’ 0.01 ‘*’ 0.05 ‘.’ 0.1 ‘ ’ 1

Residual standard error: 5.289 on 18 degrees of freedom
Multiple R-squared:  0.115,	Adjusted R-squared:  0.0658
F-statistic: 2.338 on 1 and 18 DF,  p-value: 0.1436
```
